# *HspA1B* Is a Prognostic Biomarker and Correlated With Immune Infiltrates in different subtypes of Breast Cancers

**DOI:** 10.1101/725861

**Authors:** Jian He, Hui Wang

## Abstract

**Background:** Heat shock A1B, also known as HSP70kDa protein 1B, encodes a 70kDa heat shock protein which is a member of the heat shock protein 70 family. *HspA1B* is a critical gene which related to many type of diseases by involving in the ubiquitin-proteasome pathway. However, the correlations of *HspA1B* to prognosis and tumor-infiltrating lymphocytes in different cancers remain unclear.

**Methods:** *HspA1B* expression was evaluated on the Oncomine database and Tumor Immune Estimation Resource (TIMER) site. We analyzed the influence of *HspA1B* on clinical prognosis using Kaplan-Meier plotter, the PrognoScan database and Gene Expression Profiling Interactive Analysis (GEPIA). The correlations between *HspA1B* and cancer immune infiltrates was investigated via TIMER. In addition, correlations between *HspA1B* expression and gene marker sets of immune infiltrates were analyzed by TIMER and GEPIA.

**Results:** Three cohorts (GSE9195, GSE9893, GSE3494-GPL96)) of breast cancer patients showed that high *HspA1B* expression was associated with poorer overall survival (OS), disease-specific survival (DSS), and disease-free survival (DFS). In addition, high *HspA1B* expression was significantly correlated with poor OS and progression-free survival (PFS) in bladder cancer, brain cancer and skin cancer. Moreover, *HspA1B* significantly impacts the prognosis of diverse cancers via The Cancer Genome Atlas (TCGA). *HspA1B* expression was positively correlated with infiltrating levels of CD4+ T and CD8+ T cells, macrophages, neutrophils, and dendritic cells (DCs) indifferent subtypes of Breast cancer. *HspA1B* expression showed strong correlations with diverse immune marker sets in BRCA-Luminal.

**Conclusions:** Our findings suggest that *HspA1B* is correlated with prognosis and immune infiltrating levels of, including those of CD8+ T cells, CD4+ T cells, macrophages, neutrophils, and DCs in multiple cancers, especially in colon and gastric cancer patients. In addition, *HspA1B* expression potentially contributes to regulation of tumor-associated macrophages (TAMs), DCs, T cell exhaustion and Tregs in colon and gastric cancer. These findings suggest that *HspA1B* can be used as a prognostic biomarker for determining prognosis and immune infiltration in BRCA-Luminal subtype.

## 1. INTRODUCTION

Breast Cancer is the most common and the most-studied malignancy among worldwide, and metastasis is an important biological feature that leads to a poor prognosis [1]. Researchers have made great strides in the treatment of some types of breast cancer, but the battle continues on many fronts. Immune-related mechanisms play an important role in breast cancer, and immunotherapy are considered a promising direction for the treatment of breast cancers [2, 3], whereby scientists are attempting to harness the body’s own immune system to fight and prevent malignancies [4]. Immunotherapy, such as PD-1, PD-L1 and cytotoxic T lymphocyte associated antigen 4 (CTLA4) inhibitors, has demonstrated promising anti-tumor effects in NSCLC and melanoma [4-6]. However, current immunotherapies, such as anti-CTLA4 [7-9] showed poor clinical efficacy in breast cancers, blockade of PD-1 or PD-L1 shows antitumor activity in some subsets of breast cancer patients [3]. Additionally, an increasing number of studies have found that the tumor-infiltrating lymphocytes (TIL) play an essential role in mediating response to chemotherapy and improving clinical outcomes of various cancer types [10, 11], such as tumor associated macrophages (TAMs) [12-14] and tumor-infiltrating neutrophils (TINs), they also affect the prognosis [15-18]. Therefore, there is an urgent need for the elucidation of the immunophenotypes of tumor-immune interactions and identification of novel immune-related therapeutic targets in breast cancers.

The heat shock protein family member 1B (*HspA1B*), also known as HSP70kDa protein 1B, is a protein encoding-gene located on chromosome 6, which was first reported in 1999 [19]. This intronless gene encodes a 70kDa heat shock protein which is a member of the heat shock protein 70 family [20]. Compared to the other human heat shock protein family were discovered in 1980s, the discovery of *HspA1B* was nearly twenty years later [21, 22]. *HspA1B* is expressed in many cell types and organs [26-28], it is a stress-inducible molecular chaperone with key roles that involve polypeptides refolding and degradation. It is also involved in the ubiquitin-proteasome pathway through interaction with the AU-rich element RNA-binding protein 1. The gene is located in the major histocompatibility complex class III region, in a cluster with two closely related genes which encode similar proteins [23, 24]. In conjunction with other heat shock proteins (eg. Hsp90), Hsp70 could form a complex to stabilize existing proteins against aggregation and mediates the folding of newly translated proteins in the cytosol and in organelles [25-27]. Moreover, this cochaperones complex can interact within the ligand binding domain of the glucocorticoid receptor (GR) to stabilize a specific conformational [28, 29]. Unlike other steroid hormone receptors, glucocorticoid receptor (GR) is not considered an oncogene, but it paly important role in various types of cancer. Glucocorticoids have the potential to play multiple roles in the regulation of breast cancers including their control of cellular differentiation, apoptosis and proliferation [30]. Moreover, glucocorticoids (GCs) work through GR to arrest growth and induce apoptosis in lymphoid tissue to contribute tumor invasion and metastasis [31]. All these imply that Hsp70 members play an important role in cancer. Previous studies indicate that *Hsp70A1A* is involved in pancreatic cancer and hepatocellular carcinoma [32, 33], but the relationship between Hsp70A1B and tumor progression and the mechanism which involved in is still not well defined.

Our previous research found that Single cell RNA sequencing of T cells confirmed that *HspA1B* was up regulated in activated CD8+ T and Treg cells and represses CD8+ T cell functions in vitro. These findings suggest that *HspA1B* has multifaceted functional roles in Treg cells and tumor infiltrating lymphocytes. However, the underlying functions and mechanisms of *HspA1B* in tumor progression and tumor immunology is still unclear.

In this present study, we comprehensively analyzed *HspA1B* expression and correlation with prognosis of cancer patients in databases such as Oncomine, PrognoScan, and Kaplan-Meier plotter. Moreover, we investigated the correlation of *HspA1B* with tumor-infiltrating immune cells in the different tumor microenvironments via Tumor Immune Estimation Resource (TIMER). The findings in this report shed light on the important role of *HspA1B* in colorectal and gastric cancers as well as provide a potential relationship and an underlying mechanism between *HspA1B* and tumor-immune interactions.

## 2. MATERIALS AND METHODS

### 2.1 Oncomine Database Analysis

The expression level of the *HspA1B* gene in various types of cancers was identified in the Oncomine database (https://www.oncomine.org/resource/login.html) [34]. The threshold was determined according to the following values: *P*-value of 0.0001, fold change of 1.5, and gene ranking top 5%.

### 2.2 PrognoScan Database Analysis

The correlation between *HspA1B* expression and survival in various types of cancers was analyzed by the PrognoScan (http://dna00.bio.kyutech.ac.jp/PrognoScan/) [35] and GEPIA database (http://gepia.cancer-pku.cn/) [36]. PrognoScan and GEPIA searches for relationships between gene expression and patient prognosis, such as overall survival (OS) and disease-free survival (DFS), across a large collection of publicly available cancer microarray datasets. The threshold was adjusted to a Cox *P*-value < 0.05.

### 2.3 Kaplan-Meier Plotter Database Analysis

Kaplan-Meier plotter is capable to assess the effect of 54,675 genes on survival using 10,461 cancer samples in 21 cancer types. These datasets include 6,234 breast, 2,190 ovarian, 3,452 lung, and 1,440 gastric cancer samples with a mean follow-up of 69, 40, 49, and 33 months, respectively. The correlation between *HspA1b* expression and survival in breast, ovarian, lung and gastric cancers was analyzed by Kaplan-Meier plotter (http://kmplot.com/analysis/) [37]. The hazard ratio (HR) with 95% confidence intervals and log-rank P-value were also computed.

### 2.4 TIMER Database Analysis

TIMER is a comprehensive resource for systematic analysis of immune infiltrates across diverse cancer types (https://cistrome.shinyapps.io/timer/) which applies a deconvolution previously published statistical method to infer the abundance of tumor-infiltrating immune cells (TIICs) from gene expression profiles [38, 39]. The TIMER database includes 10,897 samples across 32 cancer types from The Cancer Genome Atlas (TCGA) to estimate the abundance of immune infiltrates. We analyzed *HspA1B* expression in different types of cancer and the correlation of *HspA1B* expression with the abundance of immune infiltrates, including B cells, CD4+ T cells, CD8+ T cells, neutrophils, macrophages, and dendritic cells, via gene modules. Gene expression levels against tumor purity is displayed on the left-most panel [40, 41]. In addition, correlations between *HspA1B* expression and gene markers of tumor-infiltrating immune cells were explored via correlation modules. The gene markers of tumor-infiltrating immune cells included markers of CD8+ T cells, T cells (general), B cells, monocytes, TAMs, M1 macrophages, M2 macrophages, neutrophils, natural killer (NK) cells, dendritic cells (DCs), T-helper 1 (Th1) cells, T-helper 2 (Th2) cells, follicular helper T (Tfh) cells, T-helper 17 (Th17) cells, Tregs, and exhausted T cells. These gene markers are referenced in prior studies [42, 43]. The correlation module generated the expression scatter plots between a pair of userdefined genes in a given cancer type, together with the Spearman’s correlation and the estimated statistical significance. *HspA1B* was used for the x-axis with gene symbols, and related marker genes are represented on the y-axis as gene symbols. The gene expression level was displayed with log2 RSEM.

### 2.5 Gene Correlation Analysis in GEPIA

The online database Gene Expression Profiling Interactive Analysis 2 (GEPIA2) (http://gepia.cancer-pku.cn/index.html) was used to further confirm the significantly correlated genes. GEPIA 2 [36] is an interactive web server is a source for gene expression analysis based on 9,736 tumor and 8,587 normal samples from the TCGA and the GTEx databases.

Featuring 198619 isoforms and 84 cancer subtypes, GEPIA2 has extended gene expression quantification from the gene level to the transcript level, and supports analysis of a specific cancer subtype, and comparison between subtypes [36]. GEPIA was used to generate survival curves, including OS and DFS, based on gene expression with the log-rank test and the Mantel-Cox test in 33 different types of cancer. Gene expression correlation analysis was performed for given sets of TCGA expression data. The Spearman method was used to determine the correlation coefficient. LAYN was used for the x-axis, and other genes of interest are represented on the y-axis. The tumor and normal tissue datasets were used for analysis.

### 2.6 Statistical Analysis

Survival curves were generated by the PrognoScan and Kaplan-Meier plots. The results generated in Oncomine are displayed with P-values, fold changes, and ranks. The results of Kaplan-Meier plots, PrognoScan, and GEPIA are displayed with HR and P or Cox P-values from a log-rank test. The correlation of gene expression was evaluated by Spearman’s correlation and statistical significance, and the strength of the correlation was determined using the following guide for the absolute value: 0.00–0.19 “very weak,” 0.20–0.39 “weak,” 0.40–0.59 “moderate,” 0.60–0.79 “strong,” 0.80–1.0 “very strong.” P-values <0.05 were considered statistically significant.

## 3. RESULTS

### 3.1 The mRNA Expression Levels of HspA1b in Different Types of Human Cancers

To determine differences of *HspA1b* expression in tumor and normal tissues, the *HspA1B* mRNA levels in different tumors and normal tissues of multiple cancer types were analyzed using the Oncomine database. This analysis revealed that the *HspA1B* expression was higher in Gastric Cancer, Head and neck cancer, Lymphoma, Melanoma, and Pancreatic Cancer compared to the normal tissues and was higher in Breast Cancer, Colorectal Cancer, Esophageal Cancer, Head and neck cancer, Lung Cancer, Ovarian Cancer, Pancreatic Cancer and Sarcoma in tumor tissues (Figure 1A). In addition, lower expression was observed in Lung Cancer, Melanoma, Sarcoma, kidney Cancer and Breast Cancer in some datasets. The detailed results of *HspA1b* expression in different cancer types is summarized in **Supplementary Table 1**. To further evaluate *HspA1B* expression in human cancers, we examined *HspA1B* expression using the RNA-seq data of multiple malignancies in TCGA. The differential expression between the tumor and adjacent normal tissues for *HspA1B* across all TCGA tumors is shown in Figure 1B. *HspA1B* expression was significantly higher in **BRCA (breast invasive carcinoma)**, CHOL (cholangiocarcinoma), COAD (colon adenocarcinoma), ESCA (Esophageal carcinoma), LIHC (Liver hepatocellular carcinoma), LUAD (Lung adenocarcinoma), **LUSC(Lung squamous cell carcinoma)**, PRAD (Prostate adenocarcinoma), READ(Rectum adenocarcinoma), STAD (Stomach adenocarcinoma), UCEC (Uterine Corpus Endometrial Carcinoma), but lower expression in HNSC (head and neck cancer), KIRC(Kidney renal clear cell carcinoma), THCA (Thyroid carcinoma) compared with adjacent normal tissues.

**FIGURE 1.**
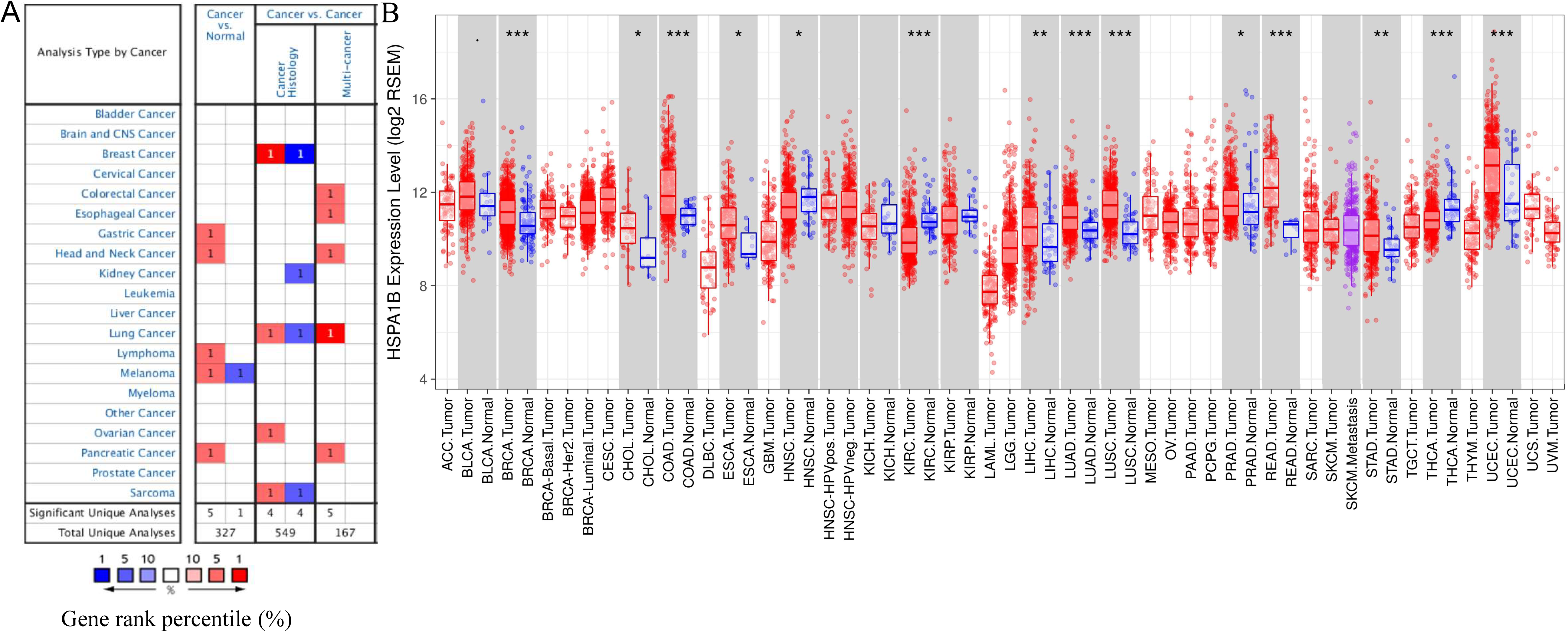
HspA1B mRNA expression levels in different types of human cancers. (A) Increased or decreased HspA1B in data sets of different cancers compared with normal tissues in the Oncomine database. Cell color is determined by the best gene rank percentile for the analyses within the cell. (B) Human HspA1B expression levels in different tumor types from TCGA database. P-value Significant Codes: 0 ≤ *** < 0.001 ≤ ** < 0.01 ≤ * < 0.05 ≤ . < 0.1

### 3.2 Prognostic Potential of *HspA1B* in Cancers

We investigated whether *HspA1B* expression was correlated with prognosis in cancer patients. The impact of *HspA1B* expression to survival rates was evaluated using the PrognoScan. The relationships between *HspA1B* expression and prognosis of different cancers in PrognoScan are shown in Supplementary Table 2. Notably, *HspA1B* expression significantly impacts OS in 4 type cancers, including Bladder Cancer, Brain cancer, **Breast cancer** and Skin cancer Supplementary Table 2. Three cohorts (GSE9195, GSE9893, GSE3494-GPL96) at different stages of Breast cancer and showed that high *HspA1b* expression was marginally associated with poorer prognosis (DMFS HR = 2.83, 95% CI = 1.19 to 6.72, Cox P = 0.018; OS=1.29, 95%CI=1.12 to 1.49, Cox P = 0.000478; DPS=1.62, 95% CI = 1.03 to 2.54, Cox P = 0.035). Therefore, it is conceivable that high *HspA1b* expression is an independent risk factor and leads to a poor prognosis in Breast cancer patients.

To further examine the prognostic potential of *HspA1B* in different cancers, Kaplan-Meier plotter database was used to evaluate the *HspA1B* prognostic value based on RNA-Seq and Affymetrix microarrays. Interestingly, the poor prognosis in Liver cancer, Kidney renal papillary cell carcinoma, **Lung adenocarcinoma**, Sarcoma, Thymoma was shown to correlate with higher *HspA1B* expression (Figures 2A–E and **Supplementary Figure 1**). In addition to RNA-Seq of *HspA1B* in the Kaplan-Meier plotter databases, the microarray analysis data also used to analyze the prognostic potential of *HspA1B* in different cancers (Breast Cancer, Gastric Cancer, Lung Cancer, Ovarian Cancer) via Kaplan-Meier plotter. The poor prognosis in Breast Cancer while a better prognosis in Gastric Cancer was shown to correlate with higher *HspA1B* expression (Figures 2H,I and **Supplementary Figure 1**) In addition to microarray analysis and RNA-Seq dataset of *HspA1b* in the PrognoScan and Kaplan-Meier plotter databases, the RNA sequencing data in TCGA were also used to analyze the prognostic potential of *HspA1b* in different cancers via GEPIA. We analyzed relationships between *HspA1b* expression and prognostic values in 33 types of cancer

**FIGURE 2.**
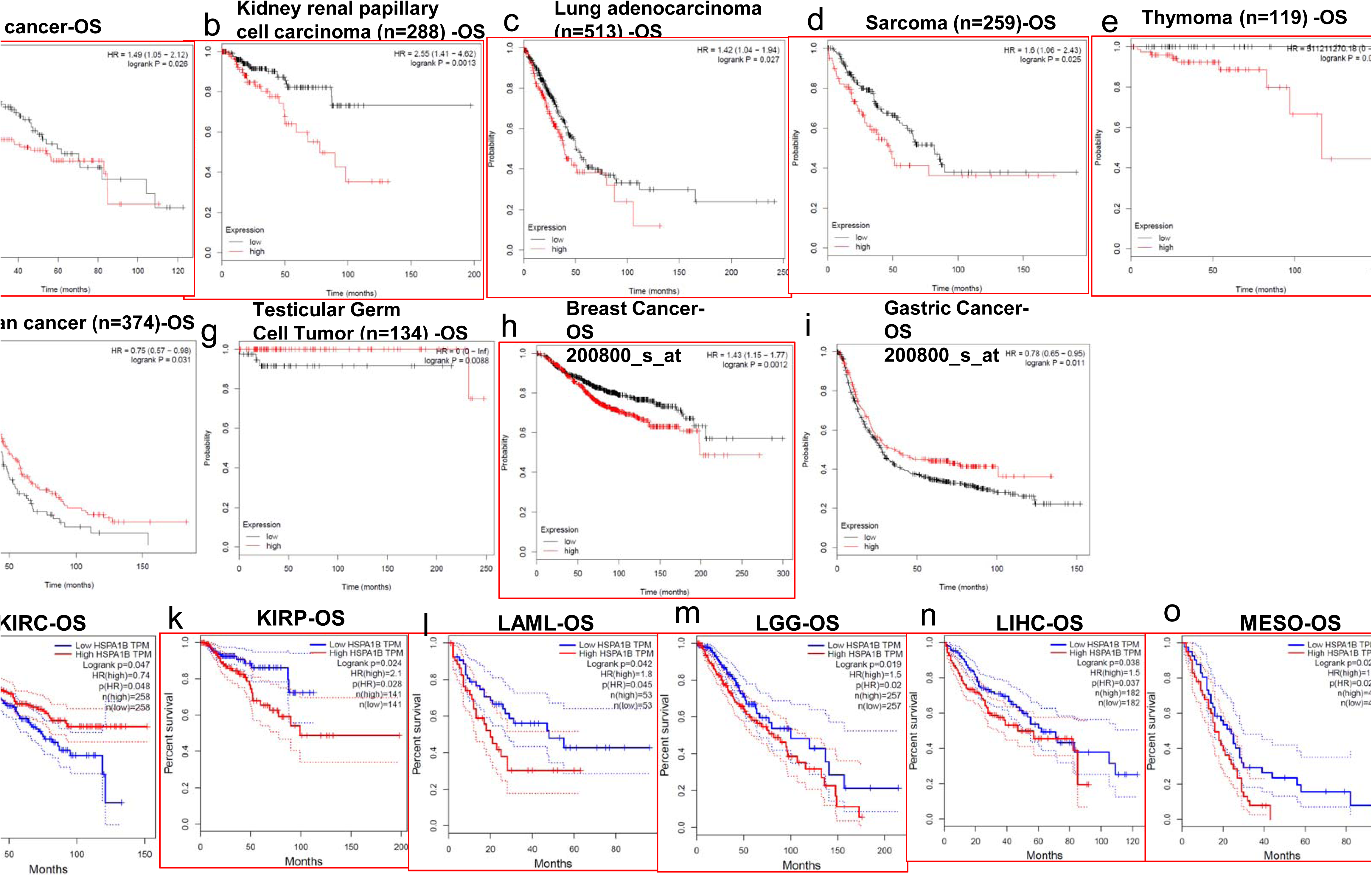
(a-e, h) The poor prognosis in Liver cancer, Kidney renal papillary cell carcinoma, Lung adenocarcinoma, Sarcoma, Thymoma, **Breast Cancer** were shown to correlate with higher *HspA1b* expression. (j–o) *HspA1b* expression significantly impacts prognosis in 6 type cancers, including KIRP(Kidney renal papillary cell carcinoma), LAML(Acute Myeloid Leukemia), LGG (Brain Lower Grade Glioma), LIHC, MESO (Mesothelioma) and SKCM and shows a better OS in KIRC(Kidney renal clear cell carcinoma)

*HspA1B* expression significantly impacts prognosis in 6 type cancers, including KIRP(Kidney renal papillary cell carcinoma), LAML(Acute Myeloid Leukemia), LGG (Brain Lower Grade Glioma), LIHC, MESO (Mesothelioma) and SKCM and shows a better OS in KIRC(Kidney renal clear cell carcinoma) (Figures 2 j–o). High *HspA1B* expression levels were associated with poorer prognosis of DFS in SKCM but has less influence on OS. These results confirmed the prognostic value of *HspA1B* in some specific types of cancers and that increased and decreased *HspA1B* expression have different prognostic value depending on the type of cancer.

### 3.3 High *HspA1B* Expression Impacts the Prognosis of Liver Cancer, Ovarian Cancer and Breast Cancer in Patients

To better understand the relevance and underlying mechanisms of *HspA1B* expression in cancer, we investigated the relationship between the *HspA1B* expression and clinical characteristics of liver cancer patients in the Kaplan-Meier plotter databases. Overexpression of *HspA1B* was associated with worse OS and PFS in female patients, race as well as different types of classification and differentiation (P < 0.05). Specifically, high *HspA1B* mRNA expression was correlated with worse OS in stage 3 to 4 and worse PFS in stage 1 to 2 of **Liver** cancer patients respectively but was not associated with OS and PFS of other stages. **(Table 1).** Since *HspA1B* showed gender correlation, we checked the female-specific cancers in the database (**SI-Fig1)**. We found that high *HspA1B* Expression was associated with worse OS and PFS in different Intrinsic subtype of breast cancer. High *HspA1B* Expression was associated with poor OS and PFS in Luminal A subtype, but no significant difference in luminal B and HER2+ subtype. High *HspA1B* mRNA expression was correlated with worse OS and PFS in Endometrioid type and worse PFS in Serous type of Ovarian cancer patients respectively but was not associated with OS and PFS of stages, grades and TP53 mutation**(Table 2).. Supplementary Figure 2.**

**TABLE 1.** Correlation of HspA1b mRNA expression and clinical prognosis in **liver cancer** with different clinicopathological factors by Kaplan-Meier plotter.

| Clinicopathological characteristics | Overall survival (n = 364) |  |  | Progression-free survival (n = 370) PFS |  |  |
| --- | --- | --- | --- | --- | --- | --- |
|  | N | Hazard ratio | P-value | N | Hazard ratio | P-value |
| **SEX** |  |  |  |  |  |  |
| Female | 118 | 2.01(1.14-3.56) | **0.014** | 120 | 1.82(1.06-3.11) | **0.027** |
| Male | 246 | 1.53(0.95-2.48) | 0.081 | 246 | 1.47(0.99-2.16) | 0.052 |
| **Race** |  |  |  |  |  |  |
| White | 184 | 1.77 (1.1 – 2.83) | **0.016** | 183 | 1.43 (0.96 – 2.13) | 0.073 |
| Black or African American | 17 | -- | -- | 17 | -- | --- |
| Asian | 155 | 1.81 (0.93 – 3.52) | 0.077 | 155 | 1.54 (0.94 – 2.52) | 0.087 |
| **STAGE** |  |  |  |  |  |  |
| 1 | 170 | 1.57 (0.77 – 3.21) | 0.21 | 170 | 1.64 (0.86 – 3.16) | 0.13 |
| 1+2 | 257 | 1.44 (0.89 – 2.35) | 0.14 | 254 | 1.58 (1.02 – 2.46) | 0.04 |
| 2 | 83 | 1.71 (0.75 – 3.89) | 0.19 | 84 | 1.57 (0.76 – 3.28) | 0.22 |
| 2+3 | 166 | 1.58 (0.97 – 2.57) | 0.062 | 167 | 1.42 (0.92 – 2.21) | 0.11 |
| 3 | 83 | 1.75 (0.95 – 3.23) | 0.07 | 83 | 1.58 (0.85 – 2.92) | 0.14 |
| 3+4 | 87 | 1.83 (1.01 – 3.32) | 0.043 | 88 | 1.57 (0.87 – 2.85) | 0.13 |
| 4 | 4 | -- | -- | 5 | -- | -- |
| **Grade** |  |  |  |  |  |  |
| 1 | 55 | 1.86 (0.68 – 5.09) | 0.22 | 55 | 3.09 (1.37 – 6.98) | 0.0045 |
| 2 | 174 | 0.57 (0.3 – 1.08) | 0.081 | 175 | 1.36 (0.86 – 2.15) | 0.18 |
| 3 | 118 | 1.49 (0.76 – 2.91) | 0.24 | 119 | 1.6 (0.83 – 3.08) | 0.15 |
| 4 | 12 | -- | -- |  | -- | -- |
| **AJCC_T** |  |  |  |  |  |  |
| 1 | 180 | 1.57 (0.79 – 3.1) | 0.19 | 180 | 1.54 (0.9 – 2.64) | 0.12 |
| 2 | 90 | 1.93 (0.9 – 4.16) | 0.087 | 92 | 1.45 (0.83 – 2.52) | 0.18 |
| 3 | 78 | 1.77 (0.95 – 3.31) | 0.071 | 78 | 1.32 (0.72 – 2.44) | 0.37 |
| 4 | 13 | -- |  | 13 | -- |  |
| **Vascular invasion** |  |  |  |  |  |  |
| none | 203 | 1.31 (0.78 – 2.2) | 0.31 | 204 | 1.56 (0.93 – 2.62) | 0.087 |
| micro | 90 | 0.55 (0.21 – 1.46) | 0.22 | 91 | 1.54 (0.87 – 2.73) | 0.14 |
| macro | 16 | -- |  | 16 |  |  |

*Bold values indicate $P < 0.05$ .*

**TABLE 2.**
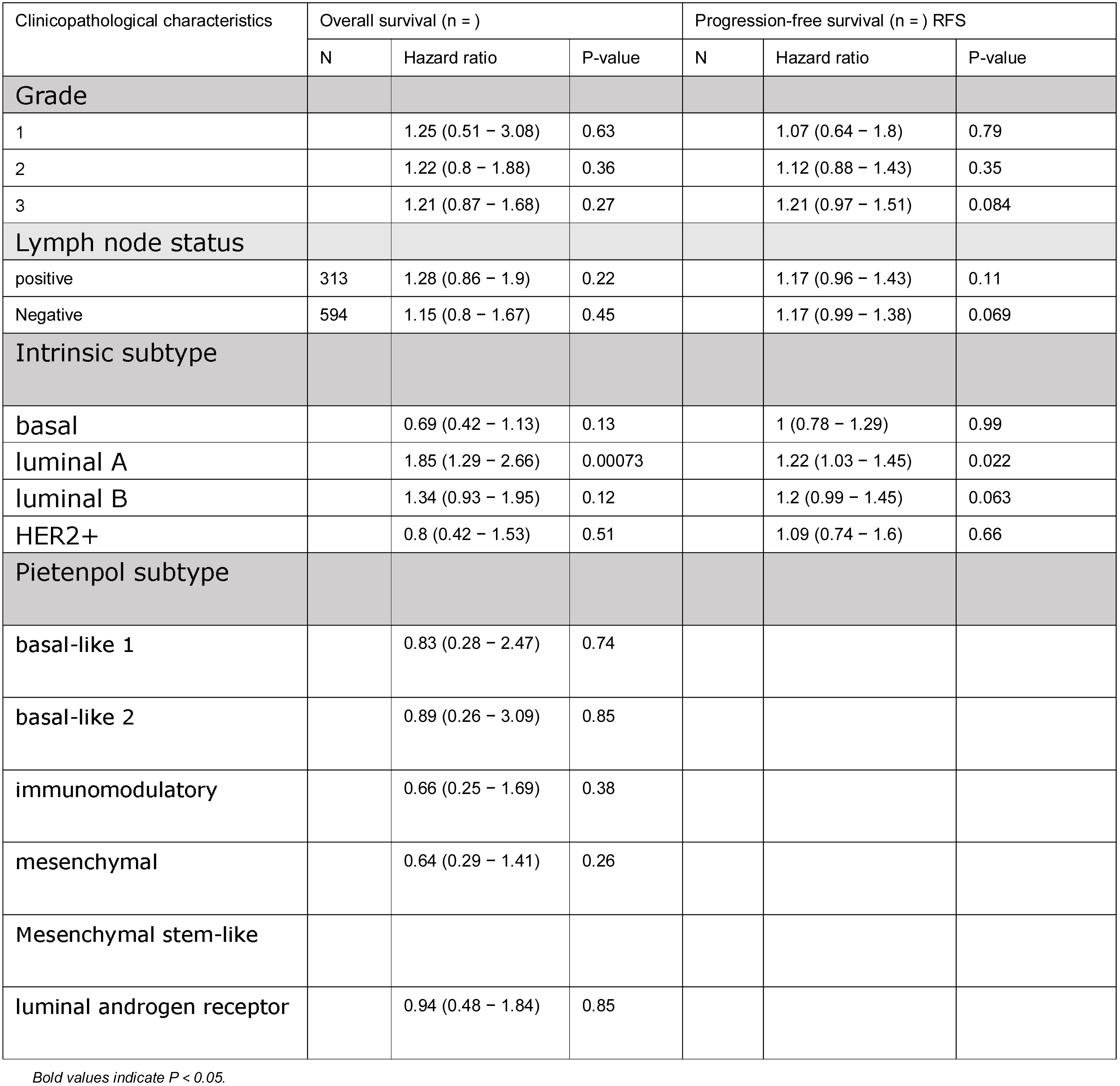
Correlation of HspA1b mRNA expression and clinical prognosis in **Breast cancer** with different clinicopathological factors by Kaplan-Meier plotter.

| Clinicopathological characteristics | Overall survival (n = ) |  |  | Progression-free survival (n = ) RFS |  |  |
| --- | --- | --- | --- | --- | --- | --- |
|  | N | Hazard ratio | P-value | N | Hazard ratio | P-value |
| **Grade** |  |  |  |  |  |  |
| 1 |  | 1.25 (0.51 – 3.08) | 0.63 |  | 1.07 (0.64 – 1.8) | 0.79 |
| 2 |  | 1.22 (0.8 – 1.88) | 0.36 |  | 1.12 (0.88 – 1.43) | 0.35 |
| 3 |  | 1.21 (0.87 – 1.68) | 0.27 |  | 1.21 (0.97 – 1.51) | 0.084 |
| **Lymph node status** |  |  |  |  |  |  |
| positive | 313 | 1.28 (0.86 – 1.9) | 0.22 |  | 1.17 (0.96 – 1.43) | 0.11 |
| Negative | 594 | 1.15 (0.8 – 1.67) | 0.45 |  | 1.17 (0.99 – 1.38) | 0.069 |
| **Intrinsic subtype** |  |  |  |  |  |  |
| basal |  | 0.69 (0.42 – 1.13) | 0.13 |  | 1 (0.78 – 1.29) | 0.99 |
| luminal A |  | 1.85 (1.29 – 2.66) | 0.00073 |  | 1.22 (1.03 – 1.45) | 0.022 |
| luminal B |  | 1.34 (0.93 – 1.95) | 0.12 |  | 1.2 (0.99 – 1.45) | 0.063 |
| HER2+ |  | 0.8 (0.42 – 1.53) | 0.51 |  | 1.09 (0.74 – 1.6) | 0.66 |
| **Pietenpol subtype** |  |  |  |  |  |  |
| basal-like 1 |  | 0.83 (0.28 – 2.47) | 0.74 |  |  |  |
| basal-like 2 |  | 0.89 (0.26 – 3.09) | 0.85 |  |  |  |
| immunomodulatory |  | 0.66 (0.25 – 1.69) | 0.38 |  |  |  |
| mesenchymal |  | 0.64 (0.29 – 1.41) | 0.26 |  |  |  |
| Mesenchymal stem-like |  |  |  |  |  |  |
| luminal androgen receptor |  | 0.94 (0.48 – 1.84) | 0.85 |  |  |  |

*Bold values indicate $P < 0.05$ .*

### 3.4 *HspA1B* Expression Is Correlated With Immune Infiltration Level in Breast Cancers

Tumor-infiltrating lymphocytes are an independent predictor of sentinel lymph node status and survival in cancers. Therefore, we investigated whether ***HspA1B*** expression was correlated with immune infiltration levels in different types of cancer. We assessed the correlations of ***HspA1B*** expression with immune infiltration levels in 34 cancer types from TIMER. The results show that ***HspA1B*** expression has significant correlations with tumor purity in 8 types (DLBC, ESCA, HNSC, LGG, LUAD, LUSC, PAAD, STAD) of cancer and significant correlations with B cell infiltration levels in 11 types of cancers (COAD, ESCA, KICH, LGG, LUAD, LUSC, PRAD, STAD, TGCT, THYM, UVM) (SI Fig3). In addition, ***HspA1B*** expression has significant correlations with infiltrating levels of CD8+ T cells in 13 types of cancer (COAD, DLBC, LUSC, PAAD, PRAD, READ, SARC, SKCM, SKCM-Primary, SKCM-Metastasis, THYM, UCS, UVM), CD4+ T cells in 11 types of cancer (COAD, ESCA, HNSC, LGG, LUAD, LUSC, PRAD, STAD, THCA, THYM, UCEC), macrophages in 14 types of cancer (BLCA, COAD, DLBC, KIRC, LGG, LUAD, LUSC, OV, PAAD, PRAD, SKCM, STAD, TGCT, THYM), neutrophils in 17 types of cancer (ACC, BLCA, COAD, HNSC, KIRC, LGG, LUAD, LUSC, PCPG, READ, SKCM, SKCM-Primary, SKCM-Metastasis, TGCT, THCA, THYM, UCEC), and dendritic cells in 17 types of cancer (BLCA, ESCA, HNSC, KIRC, LGG, LIHC, LUAD, LUSC, PAAD, PRAD, SKCM, SKCM-Primary, STAD, TGCT, THCA, THYM, UVM) (**Supplementary Figure 3**).

Given the association of *HspA1b* expression with immune infiltration level in diverse types of cancer, we next determined the distinct types of cancers in which *HspA1b* was associated with prognosis and immune infiltration. Tumor purity is an important factor that influences the analysis of immune infiltration in clinical tumor samples by genomic approaches, and TIMER and GEPIA have most of the homologous data from TCGA. Therefore, we selected the cancer types in which *HspA1b* expression levels have a significant negative correlation with tumor purity in TIMER and a significant correlation with prognosis in GEPIA. Interestingly, we found that *HspA1B* expression level correlate with poorer prognosis and high immune infiltration in KIRP, LAML, LGG, LIHC, MESO, SKCM and different Breast Cancers subtypes. *HspA1B* expression level has significant positive correlations with infiltrating levels of CD4+ T cells, macrophages, neutrophils and DCs in LGG (Fig3, SI-Fig2 AC). Similarly, there were positive correlations with infiltrating levels of CD8+ T cells, macrophages, neutrophils and DCs in SKCM (Fig3, SI-Fig2 AY). More Interestingly, the correlation with macrophages and DCs are different pattern in Primary and Metastasis subtypes of SKCM (Fig3). These findings strongly suggest that *HspA1b* plays a specific role in immune infiltration in LGG and SKCM cancers.

**FIGURE 3.**
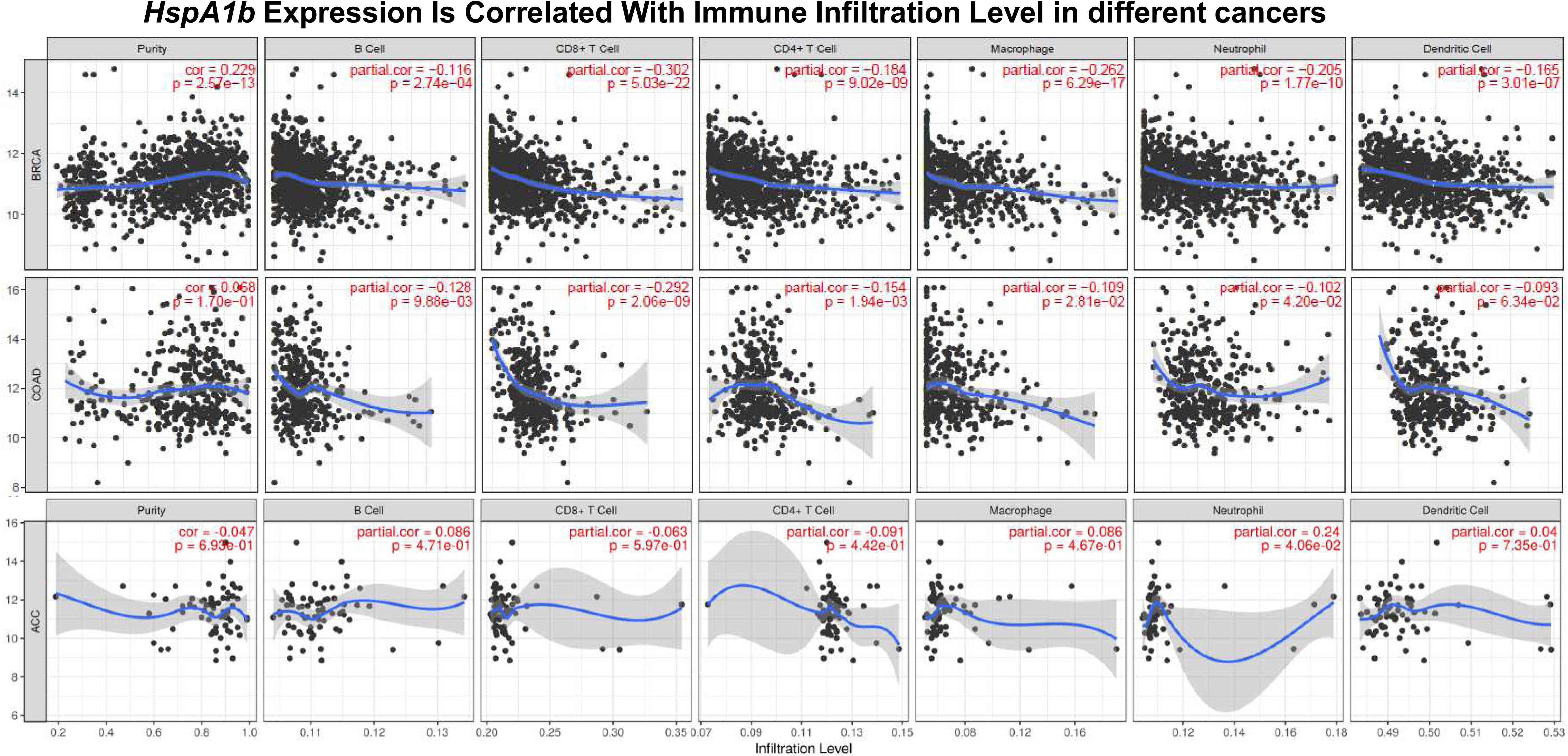
Correlation of HspA1b expression with immune infiltration level in BRCA, COAD and ACC (as control).

What attracted our attention is the *HspA1B* Expression Is Correlated With Immune Infiltration Level in different Breast Cancers subtypes (Fig 2H, Fig 4). There were positive correlations with infiltrating levels of B cells, CD8+ T cells, CD4+ T cells, macrophages, neutrophils and DCs in BRCA (Fig4 A), but the correlations with infiltrating levels of immune cells are demonstrated different pattern between Basal, Her2 and Luminal subtypes (Fig4 B-D). *HspA1B* expression level has significant positive correlations with infiltrating levels of B cells, CD8+ T cells, CD4+ T cells, macrophages, neutrophils and DCs in Luminal subtypes (Fig4 D), but none of them is correlated with Basal and Her2 subtypes of BRCA (Fig4 B, C). These findings strongly suggest that *HspA1B* plays a specific role in immune infiltration in different subtypes of BRCA.

### 3.5 Correlation Analysis Between *HspA1B* Expression and Immune Marker Sets

To investigate the relationship between *HspA1B* and the diverse immune infiltrating cells, we focused on the correlations between *HspA1B* and immune marker sets of various immune cells of LGG, SKCM and BRCA in the TIMER and GEPIA databases. We analyzed the correlations between *HspA1B* expression and immune marker genes of different immune cells, included CD8+ T cells, T cells (general), B cells, monocytes, TAMs, M1 and M2 macrophages, neutrophils, NK cells and DCs in different subtypes of BRCA, using Adrenocortical Carcinoma (AAC) as the control (Table 5). We also analyzed the different functional T cells, such as Th1 cells, Th2 cells, Tfh cells, Th17 cells, and Tregs, as well as exhausted T cells. After the correlation adjustment by purity, the results revealed the *HspA1B* expression level was significantly correlated with most immune marker sets of various immune cells and different T cells in different subtypes of BRCA. However, none of these gene markers was significantly correlated with the *HspA1B* expression level in AAC (Fig 5).

**FIGURE 4.** BRCA gene expression by TiMER.

**FIGURE 5.**
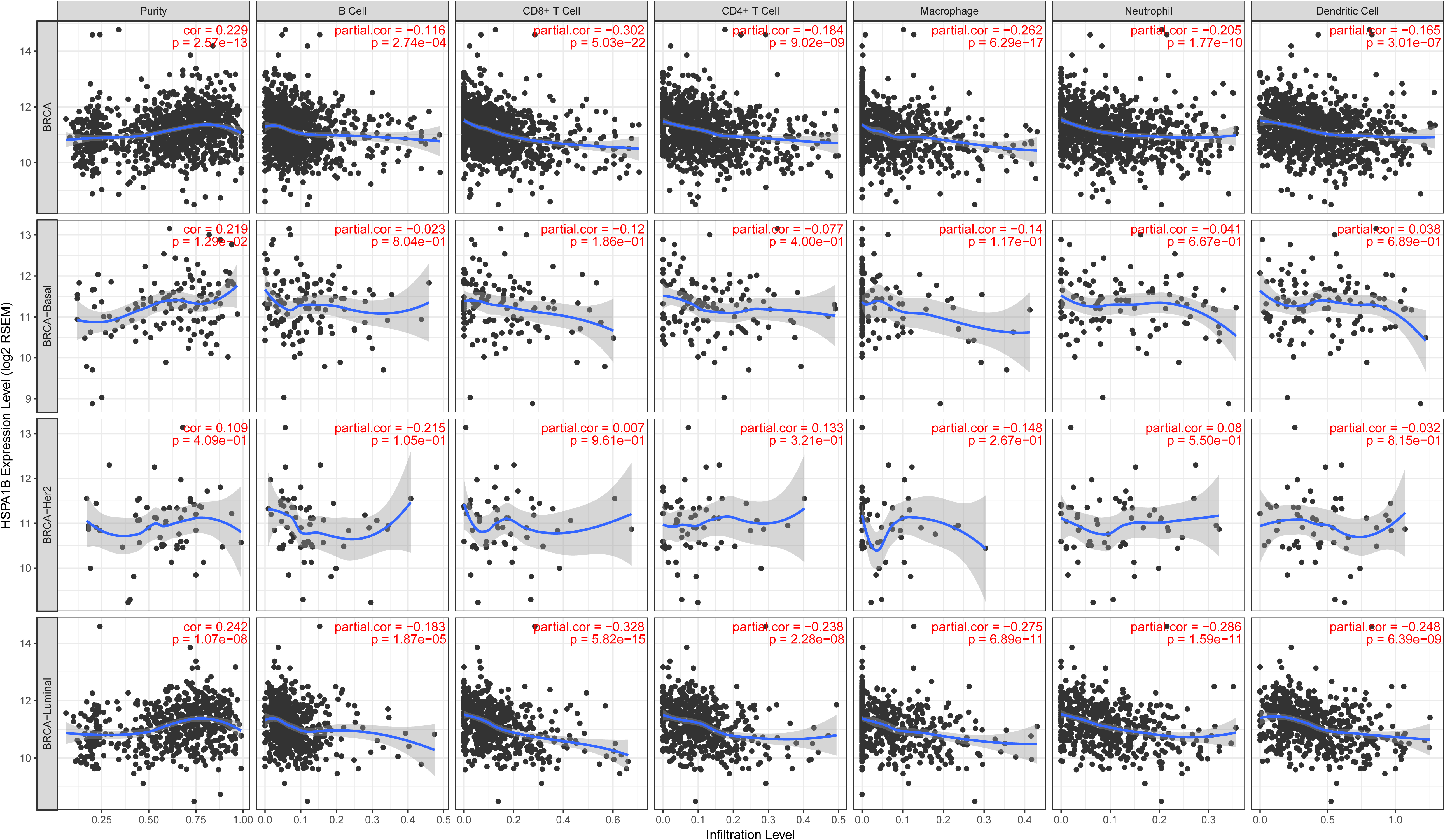
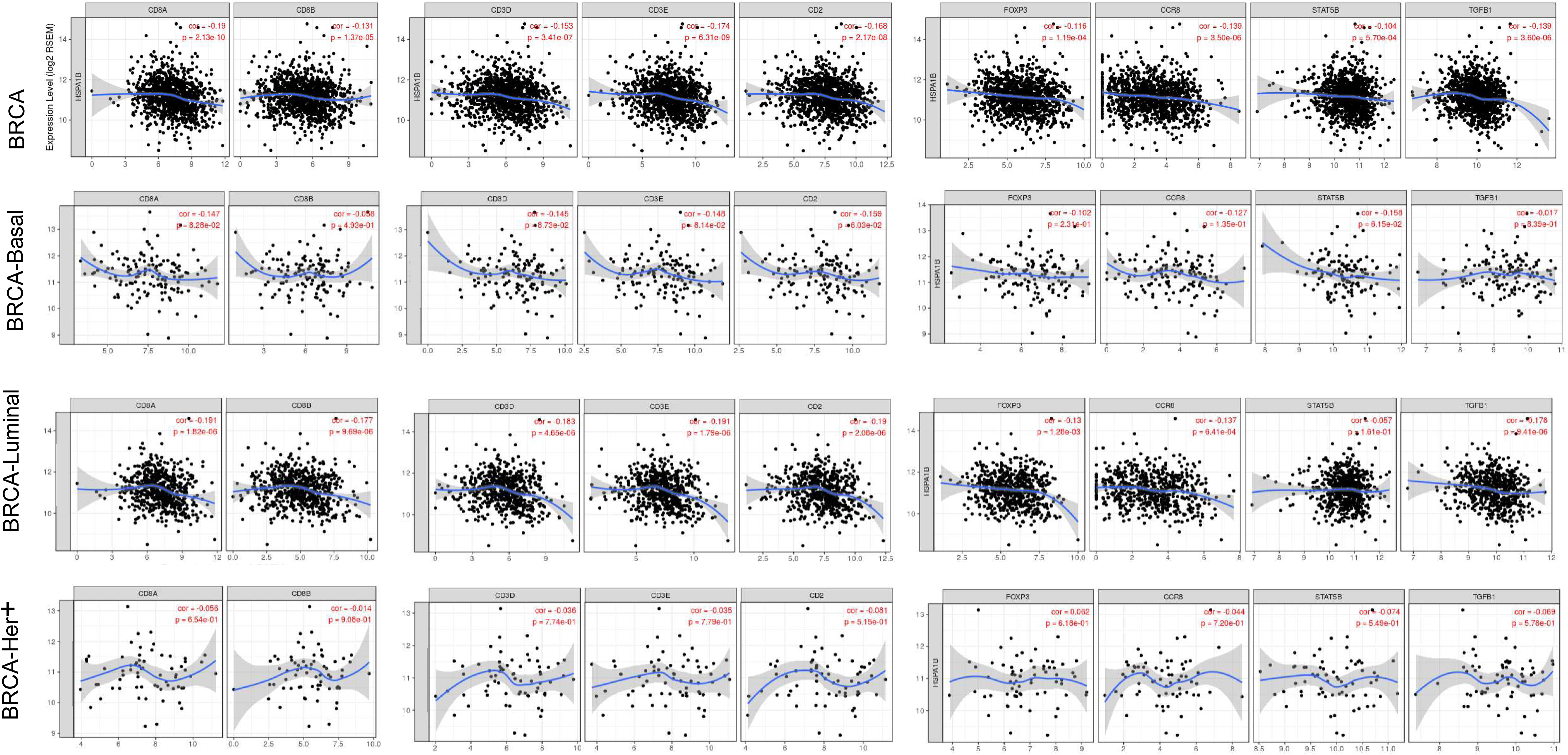
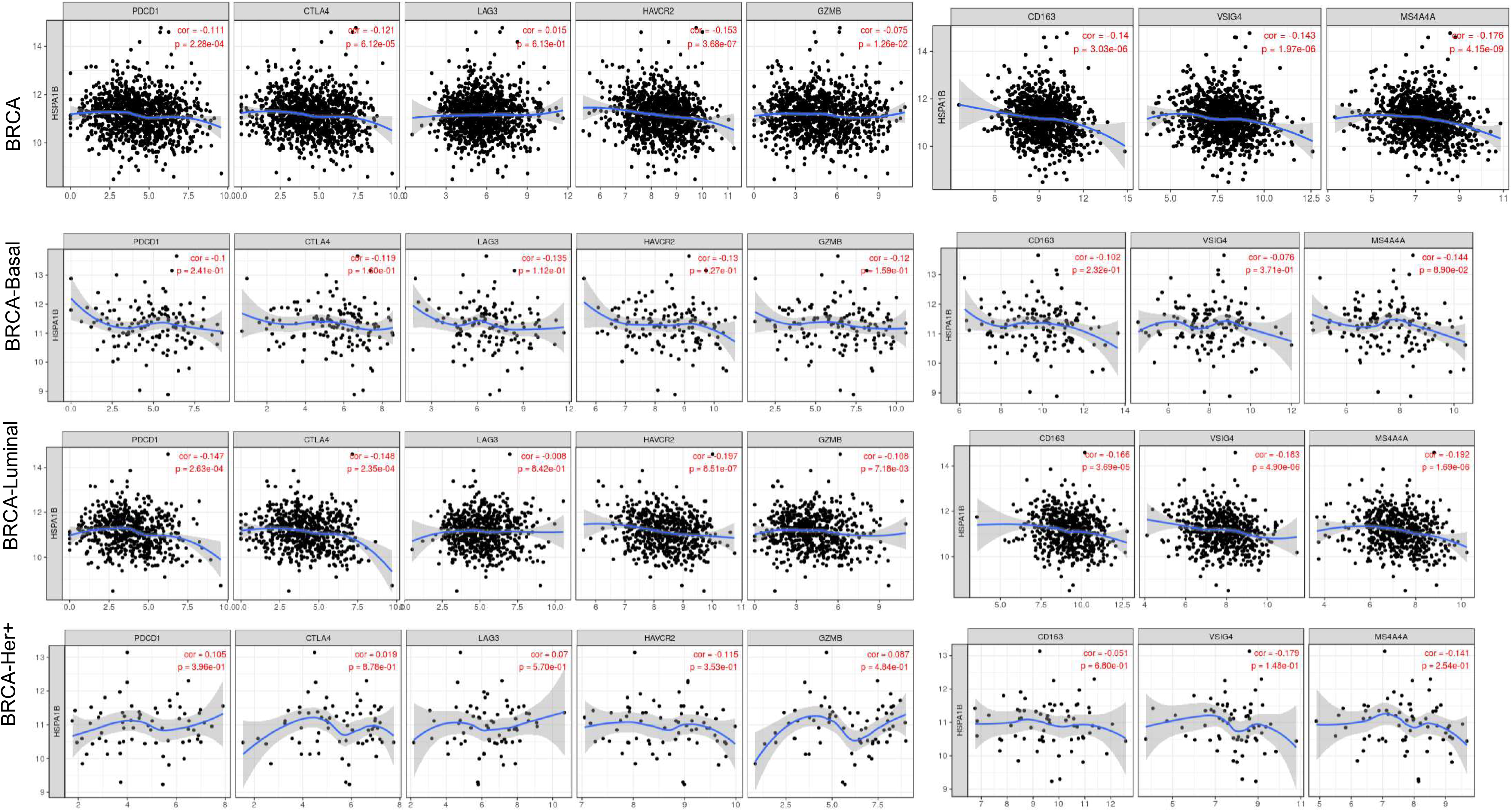
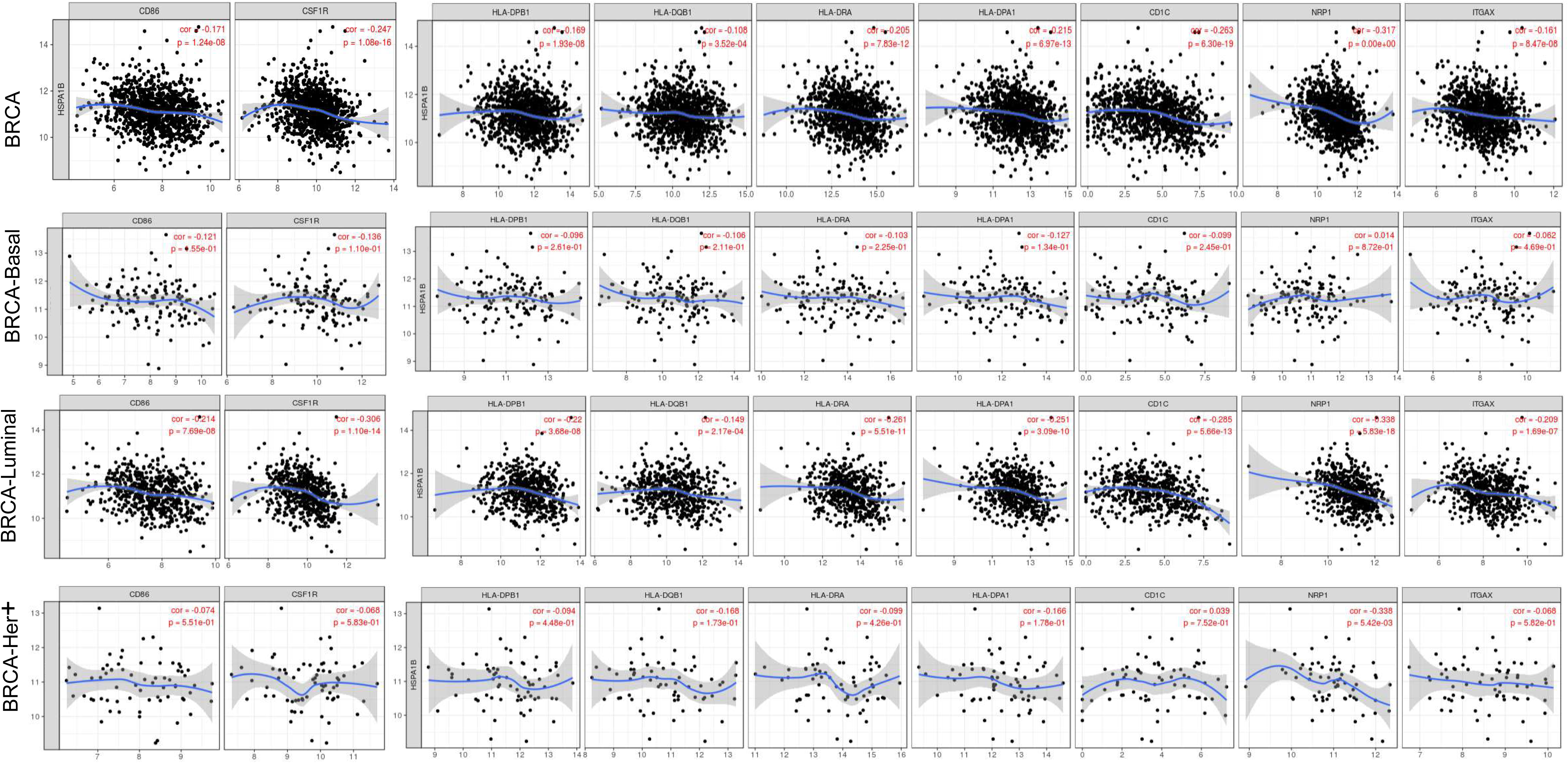
Correlation of HspA1B expression with immune infiltration level in different subtypes of Breast Cancer.

## 4. DISSCUSION

HSP70 is a stress-inducible molecular chaperone with key roles that involve polypeptides refolding and degradation. Although *HspA1B* has not been extensively studied, it is known that *HspA1B* is upregulated in activated Treg and CD8+ lymphocytes. In addition, it suppresses CD8+ functions in lung, colorectal and hepatocellular cancers. Here, we report that variations in *HspA1B* expression level correlate to prognosis in different types of cancer. High expression level of *HspA1B* correlates with a poorer prognosis in four type cancers, including Bladder Cancer, Brain cancer, Breast cancer and Skin cancer. Furthermore, our analyses show that in different subtypes of breast cancers immune infiltration levels and diverse immune marker sets are correlated with levels of *HspA1B* expression. Thus, our study provides insights in understanding the potential role of *HspA1B* in tumor immunology and its use as a cancer biomarker for different subtypes of breast cancer.

In this study, we examined the expression levels of *HspA1B* and systematic prognostic landscape in different types of cancers using independent datasets in Oncomine and 33 type cancers of TCGA data in GEPIA. The differential expression of *HspA1B* between cancer and normal tissues was observed in many types of cancer. Based on the Oncomine database, we found that *HspA1B*, compared to normal tissues, was highly expressed in Gastric, Head and neck, Pancreatic Cancer, Leukemia, Lymphoma and Melanoma, while some data sets showed that *HspA1B* has a lower level of expression in Head and neck, Lung cancer, Melanoma and Sarcoma (Figure 1A). However, analysis of the TCGA data showed that *HspA1B* expression was higher in BLCA, BRCA, CHOL, COAD, ESCA, LIHC, LUAD, LUSC, PRAD, READ, STAD, UCEC, but lower expression in HNSC, KIRC, THCA, compared with normal adjacent tissues (Figure 1B). The discrepancies in levels of *HspA1B* expression in different cancer types in different databases may be a reflection in data collection approaches and underlying mechanisms pertinent to different biological properties. Analysis of the TCGA database revealed that increased *HspA1B.* expression correlated with poor prognosis in most tumor types (KIRP, LAML, LGG, LIHC, MESO and SKCM). KIRC was exception where high levels of *HspA1B* expression showed a better prognosis.

Furthermore, analysis of data from PrognoScan showed high level of *HspA1B* expression was correlated with poor prognosis of OS in Brain cancer, Breast cancer and Skin cancer (Figures 2). Kaplan-Meier Plotter also showed a high *HspA1B* expression correlated with high hazard ratio (HR) for poor overall survival (OS) and progress free survival (PFS) of liver, Kidney renal papillary cell carcinoma, Lung adenocarcinoma, Sarcoma, Thymoma and Breast Cancer (Figures 2). In addition, high level of *HspA1B* expression was shown to be correlated with poor OS and PFS in SKCM, with poor OS and DFS, DMFS in Breast Cancer and with poor OS in two datasets (GSE4271-GPL96, MGH-glioma) of Brain Cancer. Together these findings strongly suggest that HspA1b is a prognostic biomarker in SKCM, LGG, and Breast Cancer.

Another important aspect of this study is that *HspA1B* expression is correlated with diverse immune infiltration levels in cancer, especially in SKCM, LGG, and Breast Cancer. Our results demonstrate that there is a significantly positive correlations between infiltration level of CD8+ T, CD4+ T cells, macrophages, neutrophils and DCs and *HspA1B* expression in LUSC, THYM and BRCA (Figures 3A,C). Interestingly, the correlation of the infiltration level are different in there subtypes of the breast cancer. The correlation between *HspA1B* expression and the marker genes of immune cells implicate the role of *HspA1B* in regulating tumor immunology in these types of cancers.

Firstly, gene markers of M1 macrophages such as NOS2 and IRF5 did not show correlations with *HspA1B* expression, whereas M2 macrophage markers such as CD163, VSIG4, and MS4A4A showed strong correlations in BRCA and BRCA-Luminal (Tables 2, 3). These results reveal the potential regulating role of *HspA1B* in polarization of tumor-associated macrophages (TAM), especially in BRCA-Luminal.

**TABLE 3.** Correlation of HspA1B mRNA expression and clinical prognosis in different subtypes of Breast cancer with different clinicopathological factors

| Description | Gene markers | BRCA (1093) |  | BRCA-Basal (139) |  | BRCA-Luminal (611) |  | BRCA-Her+ (67) |  |
| --- | --- | --- | --- | --- | --- | --- | --- | --- | --- |
|  |  | Cor | P | Cor | P | Cor | P | Cor | P |
| CD8+ T cell | CD8A | -0.19 | 2.13e-10 | -0.147 | 8.28e-02 | -0.191 | 1.82e-06 | -0.056 | 6.54e-01 |
|  | CD8B | -0.131 | 1.37e-05 | -0.058 | 4.39e-01 | -0.177 | 9.69e-06 | -0.014 | 9.08e-01 |
| T cell (general) | CD3D | -0.153 | 3.41e-07 | -0.145 | 8.73e-02 | -0.183 | 4.65e-06 | -0.036 | 7.74e-01 |
|  | CD3E | -0.174 | 6.31e-09 | -0.148 | 8.14e-02 | -0.191 | 1.79e-06 | -0.035 | 7.79e-01 |
|  | CD2 | -0.168 | 2.17e-08 | -0.159 | 6.03e-02 | -0.19 | 2.08e-06 | -0.081 | 5.15e-01 |
| B cell | CD19 | -0.097 | 1.25e-03 | -0.086 | 3.12e-01 | -0.1 | 1.33e-02 | 0.041 | 7.42e-01 |
|  | CD79A | -0.109 | 3.1e-04 | -0.08 | 3.45e-01 | -0.11 | 6.33e-03 | 0.03 | 8.09e-01 |
| Monocyte | CD86 | -0.171 | 1.24e-08 | -0.121 | 1.55e-01 | -0.214 | 7.69e-08 | -0.074 | 5.51e-01 |
|  | CD115 (CSF1R) | -0.247 | 1.08e-16 | -0.136 | 1.10e-01 | -0.306 | 1.1e-14 | -0.068 | 5.83e-01 |
| TAM | CCL2 | -0.051 | 9.11e-02 | -0.058 | 4.94e-01 | -0.094 | 1.95e-02 | -0.038 | 7.57e-01 |
|  | CD68 | -0.107 | 3.74e-04 | -0.094 | 2.7e-01 | -0.148 | 2.36e-04 | -0.043 | 7.29e-01 |
|  | IL10 | -0.097 | 1.23e-03 | -0.11 | 1.96e-01 | -0.112 | 5.39e-03 | -0.061 | 6.24e-01 |
| M1 Macrophage | INOS (NOS2) | -0.032 | 2.91e-01 | -0.018 | 8.37e-01 | -0.064 | 1.1e-01 | -0.014 | 9.08e-01 |
|  | IRF5 | 0.008 | 7.82e-01 | -0.058 | 4.96e-01 | -0.035 | 3.87e-01 | 0.119 | 3.35e-01 |
|  | COX2 (PTGS2) | -0.139 | 3.89e-06 | 0.13 | 1.25e-01 | -0.223 | 2.42e-08 | -0.215 | 8.05e-02 |
| M2 Macrophage | CD163 | -0.14 | 3.03e-06 | -0.102 | 2.23e-01 | -0.166 | 3.69e-05 | -0.051 | 6.8e-01 |
|  | VSIG4 | -0.143 | 1.97e-06 | -0.076 | 3.71e-01 | -0.183 | 4.9e-06 | -0.179 | 1.48e-01 |
|  | MS4A4A | -0.176 | 4.15e-09 | -0.144 | 8.9e-02 | -0.192 | 1.69e-06 | -0.141 | 2.54e-01 |
| Neutrophils | CD66b (CEACAM8) | -0.003 | 9.08e-01 | 0.046 | 5.93e-01 | -0.025 | 5.33e-01 | -0.091 | 4.64e-01 |
|  | CD11b (ITGAM) | -0.183 | 9.60e-10 | -0.065 | 4.44e-01 | -0.256 | 1.34e-10 | -0.031 | 8.04e-01 |
|  | CCR7 | -0.171 | 1.09e-08 | -0.146 | 8.62e-02 | -0.166 | 3.6e-05 | -0.152 | 2.2e-01 |
| Natural killer cell | KIR2DL1 | -0.071 | 1.88e-02 | -0.13 | 1.25e-01 | -0.094 | 1.99e-02 | 0.271 | 2.63e-02 |
|  | KIR2DL3 | -0.045 | 1.32e-01 | -0.013 | 8.77e-01 | -0.058 | 1.53e-01 | 0.168 | 1.74e-01 |
|  | KIR2DL4 | -0.037 | 2.15e-01 | -0.112 | 1.86e-01 | -0.053 | 1.93e-01 | 0.081 | 5.17e-01 |
|  | KIR3DL1 | -0.099 | 1.00e-03 | -0.093 | 2.75e-01 | -0.165 | 3.79e-05 | 0.023 | 8.55e-01 |
|  | KIR3DL2 | -0.103 | 6.00e-04 | -0.093 | 8.11e-01 | -0.124 | 1.99e-03 | -0.019 | 8.8e-01 |
|  | KIR3DL3 | -0.044 | 1.41e-01 | 0.014 | 8.74e-01 | -0.1 | 1.26e-02 | -0.003 | 9.82e-01 |
|  | KIR2DS4 | -0.047 | 1.16e-01 | -0.057 | 5.02e-01 | -0.071 | 7.56e-02 | 0.231 | 6.01e-02 |
| Dendritic cell | HLA-DPB1 | -0.169 | 1.93e-08 | -0.096 | 2.61e-01 | -0.22 | 3.68e-08 | -0.094 | 4.48e-01 |
|  | HLA-DQB1 | -0.108 | 3.52e-04 | -0.106 | 2.11e-01 | -0.149 | 2.17e-04 | -0.168 | 1.73e-01 |
|  | HLA-DRA | -0.205 | 7.83e-12 | -0.103 | 2.25e-01 | -0.261 | 5.51e-11 | -0.099 | 4.26e-01 |
|  | HLA-DPA1 | -0.215 | 6.79e-13 | -0.127 | 1.34e-01 | -0.251 | 3.09e-10 | -0.166 | 1.78e-01 |
|  | BDCA-1(CD1C) | -0.263 | 6.3e-19 | -0.099 | 2.45e-01 | -0.285 | 5.06e-13 | 0.039 | 7.52e-01 |
|  | BDCA-4(NRP1) | -0.317 | 0.00e+00 | 0.014 | 8.72e-01 | -0.338 | 5.83e-18 | -0.338 | 5.42e-03 |
|  | CD11c (ITGAX) | -0.161 | 8.47e-08 | -0.062 | 4.69e-01 | -0.209 | 1.69e-07 | -0.068 | 5.82e-01 |
| Th1 | TBX21 | -0.148 | 8.8e-07 | -0.127 | 1.35e-01 | -0.163 | 5.06e-05 | -0.014 | 9.1e-01 |
|  | SOAT1 | -0.147 | 9.8e-07 | -0.11 | 1.97e-01 | -0.245 | 8.57e-10 | -0.024 | 8.74e-01 |
|  | STAT1 | -0.09 | 2.73e-03 | -0.179 | 3.43e-02 | -0.122 | 2.5e-03 | 0.103 | 4.06e-01 |
|  | IFNG | -0.041 | 1.77e-01 | -0.059 | 4.88e-01 | -0.081 | 4.46e-02 | 0.062 | 6.19e-01 |
|  | IL13 | 0.043 | 1.52e-01 | -0.14 | 9.9e-02 | -0.002 | 9.6e-01 | 0.105 | 3.97e-01 |
| Tfh | BCL6 | -0.092 | 2.27e-03 | 0.169 | 4.59e-02 | -0.101 | 1.24e-02 | -0.113 | 3.6e-01 |
|  | IL21 | -0.069 | 2.25e-02 | -0.018 | 8.33e-01 | -0.114 | 4.72e-03 | -0.058 | 6.41e-01 |
| Th17 | STAT3 | -0.152 | 4.01e-07 | -0.081 | 3.4e-01 | -0.175 | 1.31e-05 | -0.085 | 4.92e-01 |
|  | IL17A | -0.014 | 6.47e-01 | -0.036 | 6.77e-01 | -0.067 | 9.69e-02 | -0.099 | 4.24e-01 |
| Treg | FOXP3 | -0.116 | 1.19e-04 | -0.102 | 2.31e-01 | -0.13 | 1.28e-03 | 0.062 | 6.18e-01 |
|  | CCR8 | -0.139 | 3.5e-06 | -0.127 | 1.35e-01 | -0.137 | 6.41e-04 | -0.044 | 7.2e-01 |
|  | STAT5B | -0.104 | 5.7e-04 | -0.158 | 6.15e-02 | -0.057 | 1.61e-01 | -0.074 | 5.49e-01 |
|  | TGFB1 | -0.139 | 3.6e-06 | -0.017 | 8.39e-01 | -0.178 | 9.41e-06 | -0.069 | 5.78e-01 |
| T cell exhaustion | PDCD1 (PD-1) | -0.111 | 2.28e-04 | -0.1 | 2.41e-01 | -0.147 | 2.63e-04 | 0.105 | 3.96e-01 |
|  | CTLA4 | -0.121 | 6.12e-05 | -0.119 | 1.6e-01 | -0.148 | 2.35e-04 | 0.019 | 8.78e-01 |
|  | LAG3 | 0.015 | 6.13e-01 | -0.135 | 1.12e-01 | -0.008 | 8.42e-01 | 0.07 | 5.7e-01 |
|  | HAVCR2 (TIM-3) | -0.153 | 3.68e-07 | -0.13 | 1.27e-01 | -0.197 | 8.51e-07 | -0.115 | 3.53e-01 |
|  | GZMB | -0.075 | 1.26e-02 | -0.12 | 1.59e-01 | -0.108 | 7.18e-03 | 0.087 | 4.84e-01 |

Secondly, our results indicated that *HspA1B* has the potential to activate Tregs and induce T cell exhaustion. The increase in *HspA1B* expression positively correlates with the expression of Treg and T cell exhaustion markers (FOXP3, CCR8, STAT5B, TGFB1, PD-1, CTLA4, LAG3, HAVCR2 and GZMB in BRCA and BRCA-Luminal (Table 3). HAVCR2, a crucial surface protein on exhausted T cells (38), is highly correlated with *HspA1B* expression in BRCA and BRCA-Luminal, and demonstrated different correlations pattern in another two BRCA subtypes (BRCA-Basal and BRCA-Her+). Furthermore, different correlations patterns can be found between *HspA1B* expression and the regulation of several markers of T helper cells (Th1, Th2, Tfh, and Th17) in three breast cancer subtype patterns. These correlations could be indicative of a potential mechanism where *HspA1B* regulates T cell functions in BRCA and BRCA-Luminal other than BRCA-Basal and BRCA-Her+ subtypes. Together these findings suggest that the *HspA1B* plays an important role in recruitment and regulation of immune infiltrating cells in BRCA and BRCA-Luminal.

Heat shock proteins (HSPs) are highly conserved proteins throughout evolution process, and the HSP70 is a stress-inducible molecular chaperone with key roles that involve polypeptides refolding and degradation. All members of the Hsp70 family consist of two major functional domains: an N-terminal nucleotide/ATPase binding domain (NBD) and a C-terminal substrate-binding domain (SBD) [44-46].□ Hsp70 and Hsp90 are the two main types of human heat shock proteins, both of which play an important role in tumorigenesis, development, invasion, metastasis, and so on. Hsp90 can be used as anti-cancer targets. Significant tumor suppression effects have been demonstrated by some groups by inhibiting the expression of Hsp90 in tumor cells [47-50].

The BAG-1 gene (Bcl-2 binding anti-apoptotic gene 1) is an important component of the BAG gene family and act as a anti-apoptotic gene. It encodes the multifunctional protein BAG-1 protein, which acts against a variety of cytokines and inhibits tumor cell apoptosis. The anti-apoptotic effect of BAG family proteins can be regulated by the interaction of *Hsc70/Hsp70* with the BAG domain. BAG family proteins can act as co-chaperones, indirectly regulating *Hsc70/Hsp70*-mediated protein overlap and protein degradation, and could cause elevated levels of intracellular oncoproteins [51, 52]. BAG-1 interacts with the anti-apoptotic protein Bcl-2 [52], causing tumor growth, and this effect is mediated by Hsc70/Hsp70 complex [53, 54]. The BAG-1 isoform is also highly expressed in human breast cancer [55]. It has been confirmed by immunochromatographic experiments [56] that BAG-1 was highly expressed in 147 (90%) of 160 breast cancer patients, mostly in the cytoplasm, and only a few were expressed in the nucleus. At the same time, studies have shown that [57], BAG-1 is 77.1% high in 140 patients with breast sarcoma. Multivariate analysis of the above results indicated that BAG-1 expression was significantly associated with tumor recovery and survival and survival. So all these indicated that *HspA1B* could be related to the prognosis of breast cancer closely.

Nowadays, more and more *Hsp90* inhibitors has been discovery and designed as anti-tumor reagents, among which Tanespimycin (17-AAG) has entered the phase II clinical trial stage [58-60]. Ganetespib (STA-9090) and AT13387 are also high efficiency anti-tumor chemical by inhibiting Hsp90. However, whether Hsp70 could work as an anti-tumor target is not well defined, and there are not such many inhibitors as *HSP90* under study. There is only one Hsp70 inhibitor design-VER-155008, however the IC50 is 0.5 μM, it is not a good lead for pharmaceutical production.

*HSP70* is regulated by the cell cycle and is kept in a low level in normal cells. But in tumor cells, the expression level is increased due to the stimulation by mutant or abnormal proteins [61]. Some research showed that in tumor microenvironment, Hsp70 /Hsp70 complex (HSP70 /HSP70-PCs) plays a complex and important roles. On one side, it could activate immune cells, activate cellular immunity to kill tumor cells. On the other hand, it may increase the immune escape ability of tumor cells through signaling pathways, in this way to suppress the tumor killing.

HSP70/HSP70-PCs could promote epithelial mesenchymal transition (EMT) process and the migration and invasion of tumor cells in HCC. HSP70 /HSP70-PCs could promote the proliferation of the tumor cells by activating TLR2 and TLR4 firstly and then followed by activating JNK/MAPK pathway [62-64]. Also, the expression of HSP70 is high in hepatitis C-associated HCC, suggesting that HSP70 might be a molecular target for hepatitis C-associated HCC therapy [65]. It has been reported that estrogen could induce the phosphorylation of HSP by active PI3K/Akt signaling pathway [66]. So we believe that Hsp70 might change the prognosis of breast cancer by interacting with estrogen and the immune microenvironment of cancer.

## 5. CONCLUSIONS

In this study, we provided possible mechanisms which explains why *HspA1B* expression correlates with immune infiltration and poor prognosis in some cancer types, especially in breast cancer. Therefore, interactions between *HspA1B* and TME (immune cells and estrogen) could be a potential mechanism for the correlation of *HspA1B* expression with immune infiltration and poor prognosis in BRCE-Luminal subtype.

## Author Contributions

JH conceived the project analysis the data and wrote the manuscript. HW and JV participated in discussion and language editing. HW and JV reviewed the manuscript.

## Acknowledgements

This work is supported by the Project of National Science Foundation of China (NSFC 81702730) and China Postdoctoral Science Foundation (2018M630443) and China Postdoctoral Science Special Foundation (2018T110398) for JH.

---

**Supplementary Table 1. *HspA1B* expression in Oncomine database.**

**Supplementary Table 2. Relation between HspA1b expression and patient prognosis of different cancer in Prognoscan database.**

**Supplementary Figure 1.** Correlation of *HspA1b* expression with prognostic values in diverse types of cancer. Overall survival and disease free curves comparing the high and low expression of *HspA1b* in Adrenocortical carcinoma (ACC) (A-B), Bladder Urothelial Carcinoma (BLCA) (C-D), Breast invasive carcinoma (BRCA) (E-F), Cervical squamous cell carcinoma and endocervical adenocarcinoma(CESC) (G-H), Cholangio carcinoma(CHOL) (I-J), Colon adenocarcinoma(COAD) (K-L), Lymphoid Neoplasm Diffuse Large B-cell Lymphoma(DLBC) (M-N), Esophageal carcinoma(ESCA) (O-P), Glioblastoma multiforme(GBM) (Q-R), Head and Neck squamous cell carcinoma(HNSC) (S-T), Kidney Chromophobe(KICH) (U-V), Kidney renal clear cell carcinoma(KIRC) (W-X), Kidney renal papillary cell carcinoma(KIRP) (Y-Z), Acute Myeloid Leukemia(LAML) (AA-AB), Brain Lower Grade Glioma(LGG) (AC-AD), Liver hepatocellular carcinoma(LIHC) (AE-AF), Lung adenocarcinoma(LUAD) (AG-AH), Lung squamous cell carcinoma(LUSC) (AI-AJ), Mesothelioma(MESO) (AK-AL), Ovarian serous cystadenocarcinoma(OV) (AM-AN), Pancreatic adenocarcinoma(PAAD) (AO-AP), Pheochromocytoma and Paraganglioma(PCPG) (AQ-AR), Prostate adenocarcinoma(PRAD) (AS-AT), Rectum adenocarcinoma(READ) (AU-AV), Sarcoma(SARC) (AW-AX), Skin Cutaneous Melanoma(SKCM) (AY-AZ), Stomach adenocarcinoma(STAD) (BA-BB), Testicular Germ Cell Tumors(TGCT) (BC-BD), Thyroid carcinoma(THCA) (BE-BF), Thymoma(THYM) (BG-BH), Uterine Corpus Endometrial Carcinoma(UCEC) (BI-BJ), Uterine Carcinosarcoma(UCS) (BK-BL), Uveal Melanoma(UVM) (BM-BN).

**Supplementary Figure 2** -Gene expression by TIMER.

