## Supplementary material for "*HspA1B* Is a Prognostic Biomarker and Correlated With Immune Infiltrates in different subtypes of Breast Cancers": SI-Table 1

**Supplementary Table 1. *HspA1b* expression in Oncomine database**

| **Cancer** | **Cancer type** | ***P-value*** | **Fold change** | **Rank (%)** | **Sample** | **Pubmed ID** |
| --- | --- | --- | --- | --- | --- | --- |
| Breast Cancer | Invasive Ductal Breast Carcinoma | 4.89E-11 | 1.724 | 1 | 390 | Not Published |
|  | Ductal Breast Carcinoma in Situ | 4.89E-11 | -1.724 | 1 | 3 | Not Published |
| Colorectal Cancer | Colorectal Cancer | 6.19E-6 | 2.197 | 3 | 23 | Not Published |
| Esophageal Cancer | Esophageal Cancer | 1.76E-8 | 2.083 | 2 | 25 | 22460905 |
| Gastric Cancer | Gastric Intestinal Type Adenocarcinoma | 1.53E-11 | 4.706 | 2 | 26 | 19081245 |
| Head and Neck Cancer | Head and Neck Squamous Cell Carcinoma | 1.30E-5 | 2.763 | 2 | 34 | 14676830 |
|  | Oral Cavity Squamous Cell Carcinoma | 5.63E-7 | -2.086 | 8 | 57 | 21853135 |
| Kidney Cancer | Clear Cell Renal Cell Carcinoma | 2.96E-9 | -2.058 | 5 | 184 | Not Published |
| Lung Cancer | Squamous Cell Lung Carcinoma | 1.56E-6 | 2.053 | 4 | 53 | 16273092 |
|  | Non-Small Cell Lung Carcinoma | 1.56E-6 | -2.053 | 4 | 58 | 16273092 |
|  | Lung Cancer | 1.22E-5 | 2.677 | 1 | 9 | 17339364 |
| Lymphoma | Hodgkin's Lymphoma | 1.91E-5 | 2.428 | 4 | 12 | 18794340 |
| Melanoma | Benign Melanocytic Skin Nevus | 3.06E-5 | 2.418 | 3 | 18 | 16243793 |
|  | Cutaneous Melanoma | 3.62E-5 | -3.283 | 2 | 14 | 18442402 |
| Ovarian Cancer | Ovarian Serous Adenocarcinoma | 4.46E-5 | 2.326 | 3 | 71 | 20492709 |
| Pancreatic Cancer | Pancreatic Carcinoma | 1.23E-6 | 3.304 | 3 | 36 | 19732725 |
| Sarcoma | Pleomorphic Liposarcoma | 5.90E-5 | 2.102 | 3 | 23 | 20601955 |
|  | Myxoid/Round Cell Liposarcoma | 1.03E-12 | -3.264 | 4 | 20 | 20601955 |
