## Supplementary material for "*HspA1B* Is a Prognostic Biomarker and Correlated With Immune Infiltrates in different subtypes of Breast Cancers": SI-Table 2

Supplementary Table 2. Relation between HspA1b expression and patient progonsis of different cancer in Prognoscan database.

| CANCER TYPE | DATASET | ENDPOINT | N | HR [95% CI-low CI-up] | COX P-VALUE |
|---|---|---|---|---|---|
| Bladder cancer | GSE5287 | OS | 30 | 1.13 [0.76 - 1.68] | 0.537437 |
| | GSE13507 | OS | 165 | 1.31 [0.98 - 1.76] | 0.069609 |
| | GSE13507 | DPS | 165 | 1.79 [1.07 - 2.98] | 0.0264068 |
| Blood cancer | GSE12417-GPL96 | OS | 163 | 1.10 [0.96 - 1.26] | 0.182355 |
| | GSE12417-GPL570 | OS | 79 | 1.14 [0.91 - 1.44] | 0.249759 |
| | GSE5122 | OS | 58 | 1.34 [0.84 - 2.16] | 0.218763 |
| | GSE8970 | OS | 34 | 1.49 [0.90 - 2.49] | 0.123395 |
| | GSE4475 | OS | 158 | 1.05 [0.78 - 1.42] | 0.727644 |
| | E-TABM-346 | OS | 53 | 0.83 [0.49 - 1.41] | 0.493064 |
| | E-TABM-346 | EFS | 53 | 0.79 [0.47 - 1.31] | 0.356988 |
| | GSE16131-GPL96 | OS | 180 | 1.00 [0.78 - 1.28] | 0.97588 |
| | GSE2658 | DPS | 559 | 0.74 [0.59 - 0.94] | 0.014763 |
| Brain cancer | GSE4271-GPL96 | OS | 77 | 1.39 [1.01 - 1.92] | 0.0424679 |
| | GSE7696 | OS | 70 | 1.11 [0.87 - 1.42] | 0.41109 |
| | MGH-glioma | OS | 50 | 1.09 [0.61 - 1.96] | 0.777507 |
| | MGH-glioma | OS | 50 | 1.45 [1.13 - 1.87] | 0.00402255 |
| | GSE4412-GPL96 | OS | 74 | 1.23 [0.84 - 1.79] | 0.282447 |
| | GSE16581 | OS | 67 | 1.60 [0.69 - 3.70] | 0.2722 |
| Breast cancer | GSE19615 | DMFS | 115 | 0.89 [0.40 - 1.98] | 0.769938 |
| | GSE3143 | OS | 158 | 0.94 [0.60 - 1.46] | 0.77645 |
| | GSE3143 | OS | 158 | 0.75 [0.48 - 1.18] | 0.211627 |
| | GSE7849 | DFS | 76 | 2.56 [1.17 - 5.61] | 0.0186585 |
| | GSE7849 | DFS | 76 | 2.53 [0.70 - 9.17] | 0.158348 |
| | GSE12276 | Relapse Free Survival | 204 | 1.28 [0.99 - 1.66] | 0.0614322 |
| | GSE6532-GPL570 | Relapse Free Survival | 87 | 1.00 [0.63 - 1.58] | 0.997857 |
| | GSE6532-GPL570 | DMFS | 87 | 1.00 [0.63 - 1.58] | 0.997857 |
| | GSE9195 | Relapse Free Survival | 77 | 1.89 [0.89 - 4.01] | 0.0949971 |
| | GSE9195 | DMFS | 77 | 2.83 [1.19 - 6.72] | 0.0181044 |
| | GSE12093 | DMFS | 136 | 0.98 [0.52 - 1.85] | 0.943939 |
| | GSE11121 | DMFS | 200 | 1.79 [0.93 - 3.42] | 0.0799157 |
| | GSE9893 | OS | 155 | 1.29 [1.12 - 1.49] | 0.000479016 |
| | GSE2034 | DMFS | 286 | 1.32 [0.95 - 1.82] | 0.0942784 |
| | GSE1456-GPL96 | OS | 159 | 1.44 [0.88 - 2.35] | 0.147119 |
| | GSE1456-GPL96 | Relapse Free Survival | 159 | 1.07 [0.66 - 1.75] | 0.778342 |
| | GSE1456-GPL96 | DPS | 159 | 1.52 [0.85 - 2.71] | 0.157527 |
| | GSE7378 | DFS | 54 | 1.67 [0.63 - 4.42] | 0.30159 |
| | E-TABM-158 | DMFS | 117 | 1.14 [0.74 - 1.76] | 0.541928 |
| | E-TABM-158 | OS | 117 | 1.01 [0.72 - 1.40] | 0.975536 |
| | E-TABM-158 | Relapse Free Survival | 117 | 1.01 [0.72 - 1.40] | 0.975536 |
| | E-TABM-158 | DPS | 117 | 1.15 [0.76 - 1.74] | 0.50763 |
| | GSE3494-GPL96 | DPS | 236 | 1.62 [1.03 - 2.54] | 0.0351941 |
| | GSE4922-GPL96 | DFS | 249 | 1.21 [0.85 - 1.71] | 0.284789 |
| | GSE2990 | DMFS | 125 | 1.46 [0.93 - 2.30] | 0.0964254 |
| | GSE2990 | Relapse Free Survival | 125 | 1.23 [0.88 - 1.71] | 0.22432 |
| | GSE2990 | DMFS | 54 | 0.77 [0.40 - 1.49] | 0.440647 |
| | GSE2990 | Relapse Free Survival | 62 | 0.94 [0.55 - 1.59] | 0.814162 |
| | GSE7390 | Relapse Free Survival | 198 | 1.06 [0.80 - 1.40] | 0.704012 |
| | GSE7390 | DMFS | 198 | 1.05 [0.74 - 1.49] | 0.782908 |
| | GSE7390 | OS | 198 | 1.08 [0.76 - 1.55] | 0.663504 |
| Colorectal cancer | GSE12945 | DFS | 51 | 1.13 [0.45 - 2.80] | 0.79455 |
| | GSE12945 | OS | 62 | 1.89 [0.94 - 3.81] | 0.0740171 |
| | GSE17536 | DFS | 145 | 1.16 [0.74 - 1.81] | 0.526906 |
| | GSE17536 | DPS | 177 | 0.92 [0.66 - 1.30] | 0.640755 |
| | GSE17536 | OS | 177 | 1.05 [0.78 - 1.43] | 0.73192 |
| | GSE14333 | DFS | 226 | 1.39 [0.96 - 2.01] | 0.0836341 |
| | GSE17537 | OS | 55 | 1.16 [0.71 - 1.90] | 0.548762 |
| | GSE17537 | DFS | 55 | 1.02 [0.61 - 1.71] | 0.948532 |
| | GSE17537 | DPS | 49 | 1.42 [0.74 - 2.76] | 0.294018 |
| Eye cancer | GSE22138 | DMFS | 63 | 0.91 [0.71 - 1.18] | 0.483694 |
| Head and neck cancer | GSE2837 | Relapse Free Survival | 28 | 0.97 [0.75 - 1.25] | 0.807422 |
| Lung cancer | jacob-00182-CANDF | OS | 82 | 1.16 [0.73 - 1.84] | 0.519288 |
| | HARVARD-LC | OS | 84 | 1.26 [0.85 - 1.87] | 0.242532 |
| | HARVARD-LC | OS | 84 | 1.95 [0.58 - 6.61] | 0.283689 |
| | jacob-00182-HLM | OS | 79 | 1.10 [0.79 - 1.52] | 0.584921 |
| | MICHIGAN-LC | OS | 86 | 0.91 [0.51 - 1.61] | 0.745409 |
| | jacob-00182-MSK | OS | 104 | 0.75 [0.43 - 1.30] | 0.298671 |
| | GSE31210 | OS | 204 | 1.41 [0.69 - 2.86] | 0.342437 |
| | GSE31210 | Relapse Free Survival | 204 | 1.03 [0.61 - 1.74] | 0.900636 |
| | jacob-00182-UM | OS | 178 | 1.22 [0.91 - 1.63] | 0.180117 |
| | GSE3141 | OS | 111 | 1.26 [0.90 - 1.76] | 0.183112 |
| | GSE14814 | OS | 90 | 1.03 [0.72 - 1.48] | 0.868274 |
| | GSE14814 | DPS | 90 | 1.03 [0.68 - 1.55] | 0.899362 |
| | GSE8894 | Relapse Free Survival | 138 | 1.09 [0.93 - 1.27] | 0.277438 |
| | GSE4573 | OS | 129 | 0.91 [0.57 - 1.46] | 0.70764 |
| Ovarian cancer | GSE9891 | OS | 278 | 0.99 [0.82 - 1.18] | 0.881987 |
| | DUKE-OC | OS | 133 | 0.89 [0.76 - 1.04] | 0.127315 |
| | GSE26712 | DFS | 185 | 0.98 [0.84 - 1.13] | 0.753096 |
| | GSE26712 | OS | 185 | 0.93 [0.79 - 1.10] | 0.405235 |
| | GSE14764 | OS | 80 | 0.90 [0.57 - 1.43] | 0.667811 |
| Prostate cancer | GSE16560 | OS | 281 | 1.01 [0.69 - 1.49] | 0.953527 |
| Skin cancer | GSE19234 | OS | 38 | 2.54 [1.35 - 4.78] | 0.0036724 |
| Soft tissue cancer | GSE30929 | Distant Recurrence Free Survival | 140 | 1.91 [1.39 - 2.63] | 7.45E-05 |

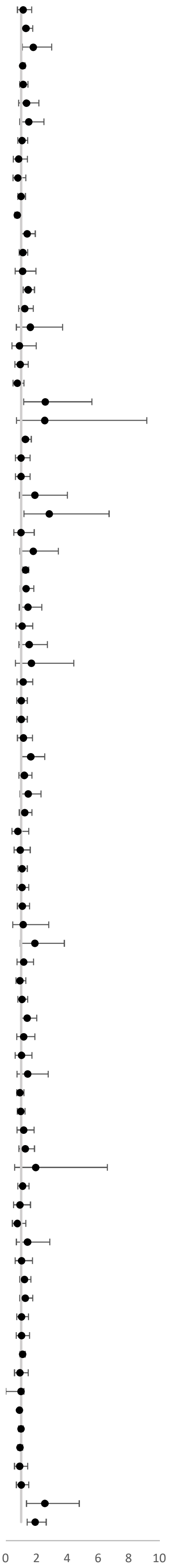
