## Supplementary material for "*HspA1B* Is a Prognostic Biomarker and Correlated With Immune Infiltrates in different subtypes of Breast Cancers": SI-Fig 1

### Slide 1
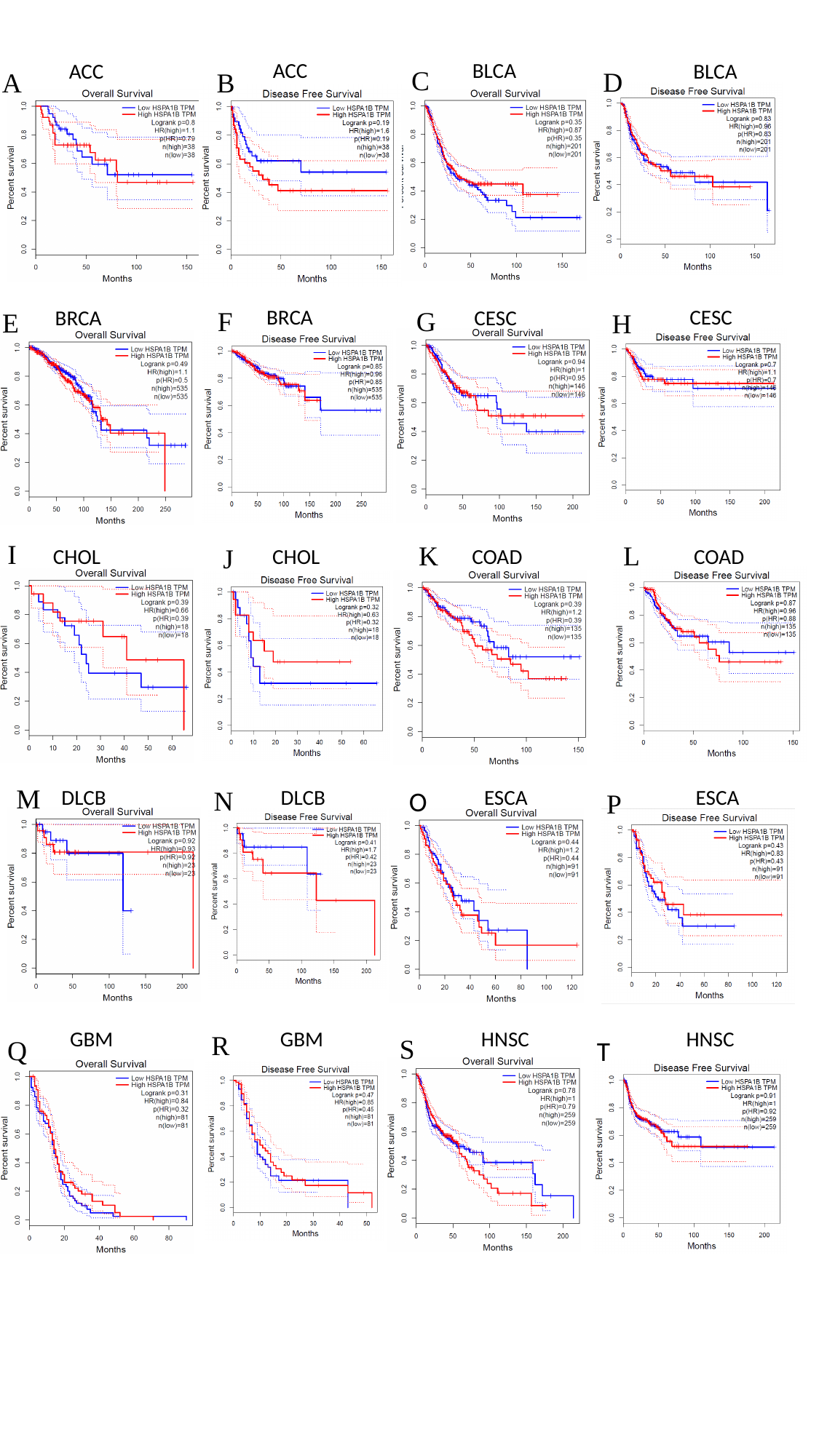

ACC
BLCA
ACC
BLCA
C
D
A
B
BRCA
CESC
CESC
BRCA
F
G
E
H
I
K
L
J
COAD
COAD
CHOL
CHOL
M
DLCB
DLCB
ESCA
ESCA
N
O
P
GBM
GBM
HNSC
HNSC
R
S
Q
T

### Slide 2
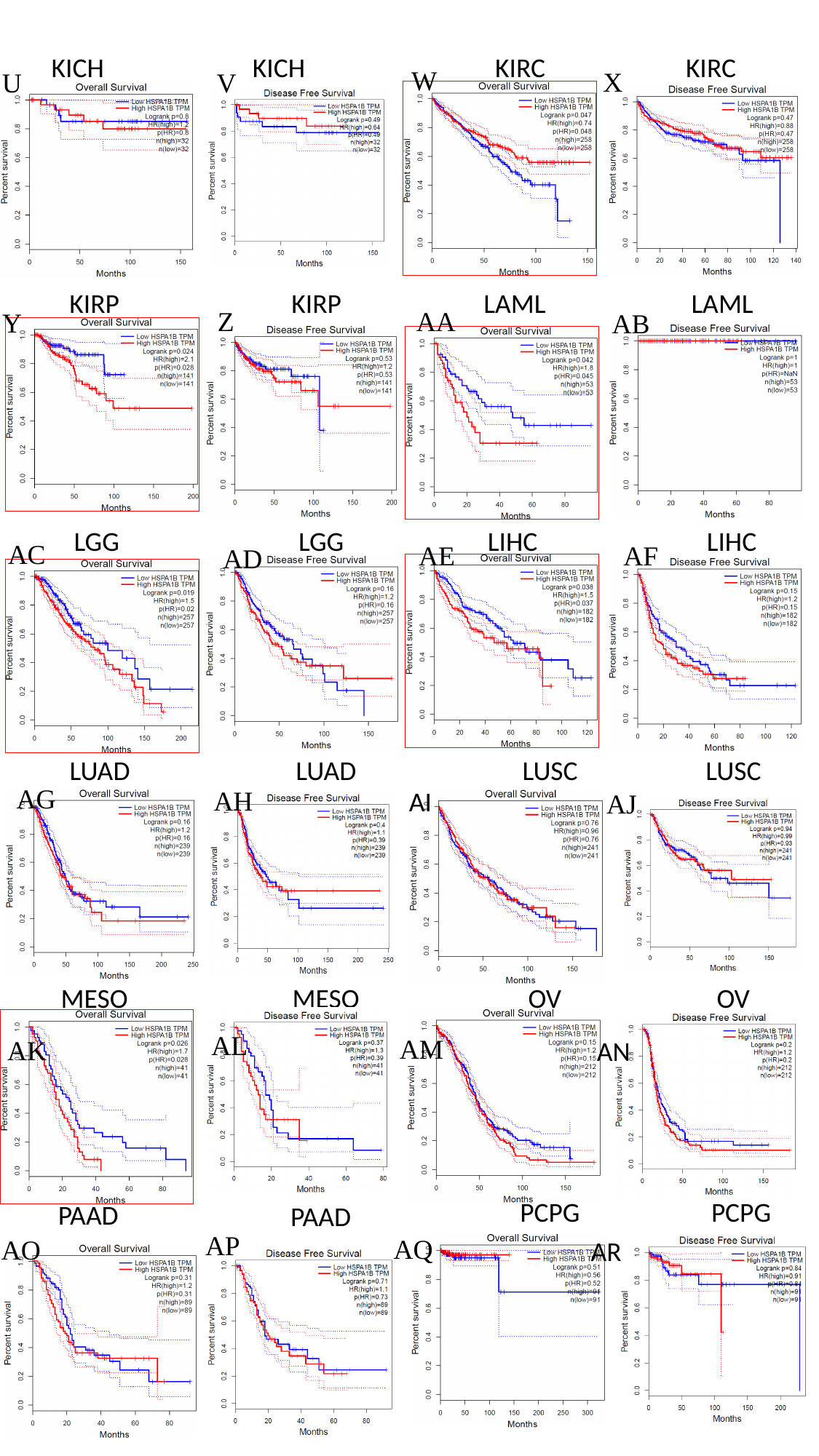

KICH
KICH
KIRC
KIRC
W
X
U
V
KIRP
KIRP
LAML
LAML
Z
AA
Y
AB
LGG
LGG
LIHC
LIHC
AC
AE
AF
AD
LUAD
LUAD
LUSC
LUSC
AG
AH
AI
AJ
MESO
MESO
OV
OV
AL
AM
AK
AN
PCPG
PCPG
PAAD
PAAD
AP
AQ
AO
AR

### Slide 3
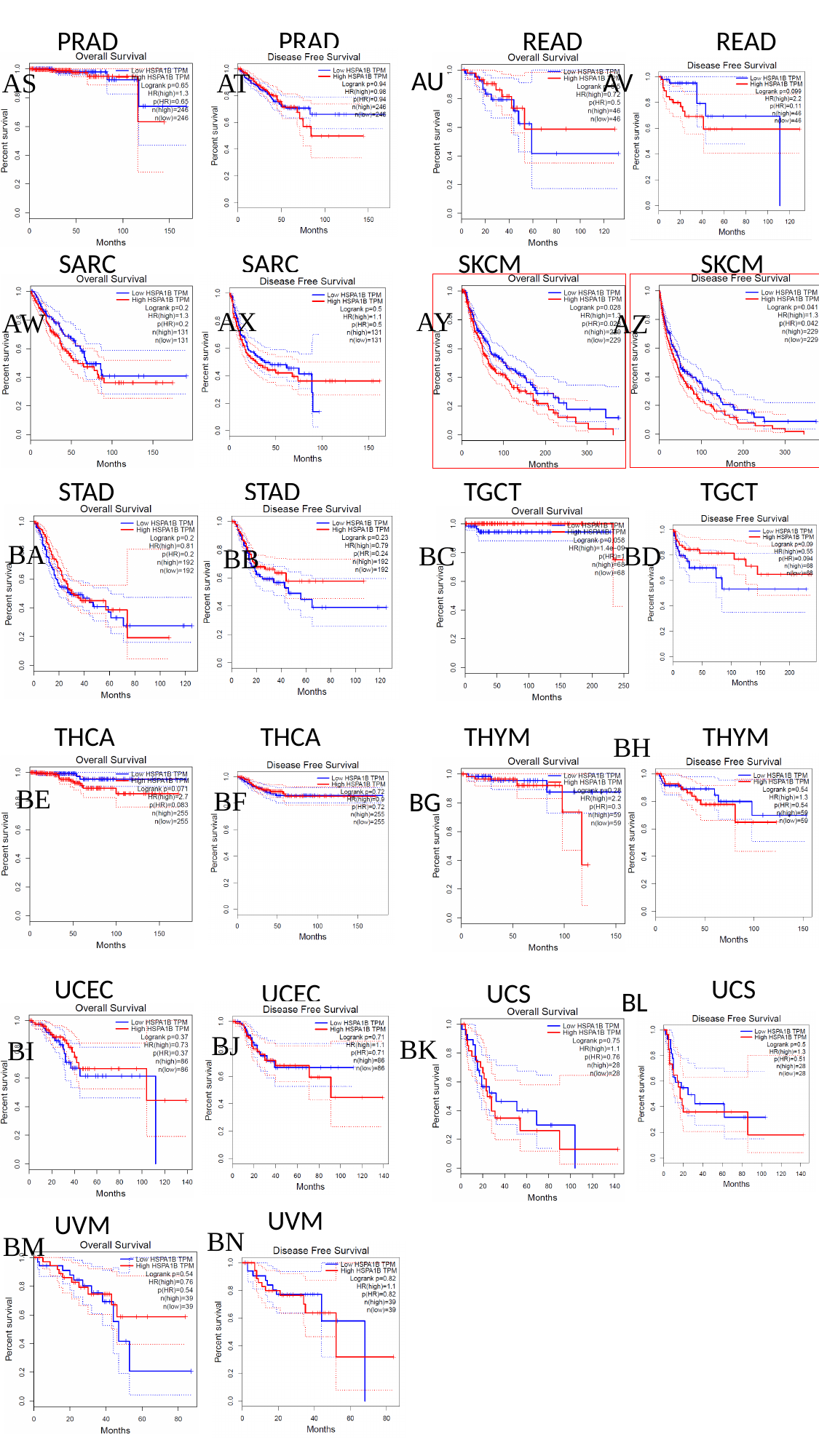

PRAD
PRAD
READ
READ
AU
AV
AS
AT
SARC
SARC
SKCM
SKCM
AX
AY
AW
AZ
STAD
STAD
TGCT
TGCT
BA
BC
BD
BB
THCA
THCA
THYM
THYM
BH
BE
BF
BG
UCEC
UCS
UCEC
UCS
BL
BJ
BK
BI
UVM
UVM
BN
BM

### Slide 4
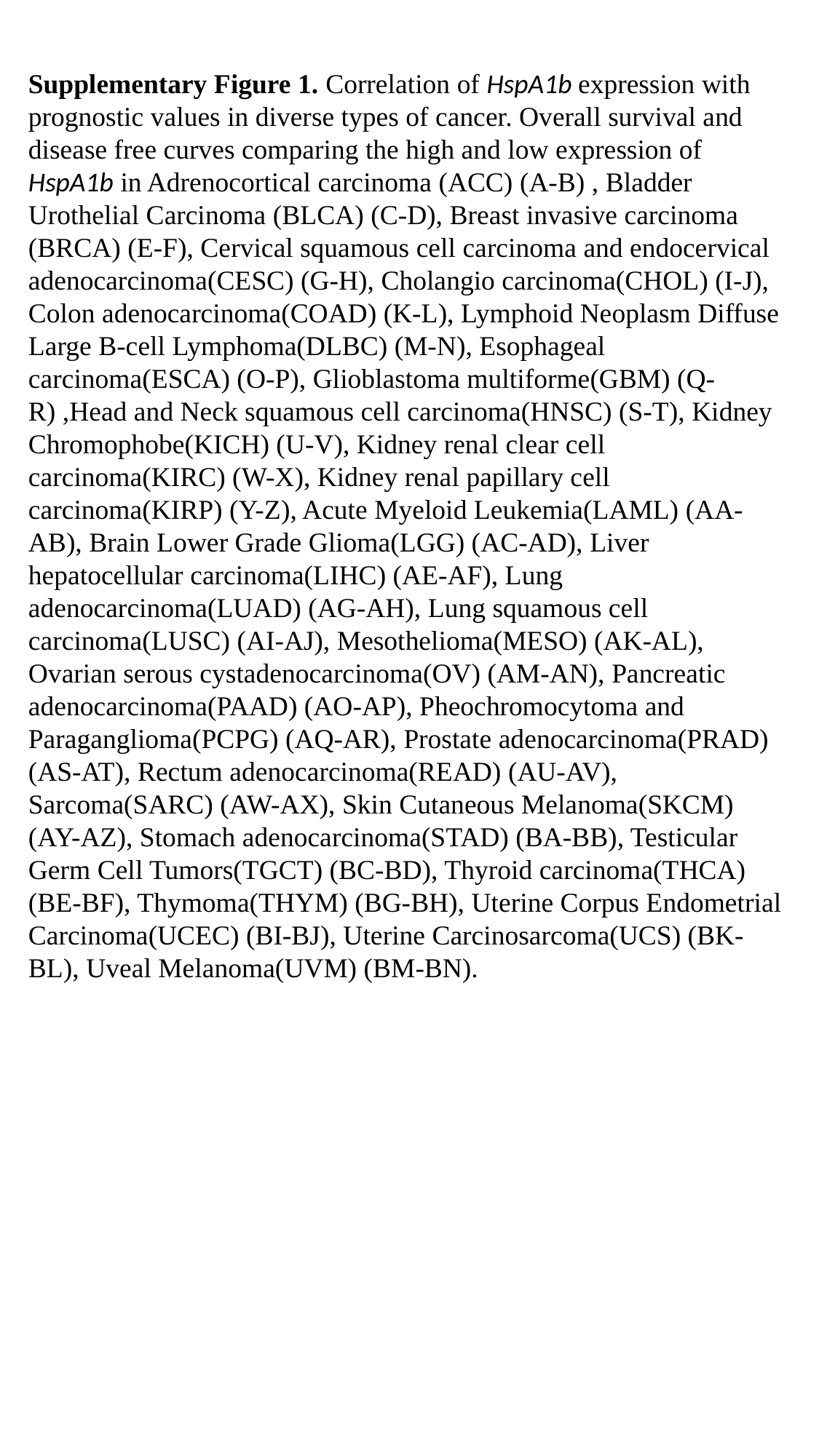

Supplementary Figure 1. Correlation of HspA1b expression with prognostic values in diverse types of cancer. Overall survival and disease free curves comparing the high and low expression of HspA1b in Adrenocortical carcinoma (ACC) (A-B) , Bladder Urothelial Carcinoma (BLCA) (C-D), Breast invasive carcinoma (BRCA) (E-F), Cervical squamous cell carcinoma and endocervical adenocarcinoma(CESC) (G-H), Cholangio carcinoma(CHOL) (I-J), Colon adenocarcinoma(COAD) (K-L), Lymphoid Neoplasm Diffuse Large B-cell Lymphoma(DLBC) (M-N), Esophageal carcinoma(ESCA) (O-P), Glioblastoma multiforme(GBM) (Q-R) ,Head and Neck squamous cell carcinoma(HNSC) (S-T), Kidney Chromophobe(KICH) (U-V), Kidney renal clear cell carcinoma(KIRC) (W-X), Kidney renal papillary cell carcinoma(KIRP) (Y-Z), Acute Myeloid Leukemia(LAML) (AA-AB), Brain Lower Grade Glioma(LGG) (AC-AD), Liver hepatocellular carcinoma(LIHC) (AE-AF), Lung adenocarcinoma(LUAD) (AG-AH), Lung squamous cell carcinoma(LUSC) (AI-AJ), Mesothelioma(MESO) (AK-AL), Ovarian serous cystadenocarcinoma(OV) (AM-AN), Pancreatic adenocarcinoma(PAAD) (AO-AP), Pheochromocytoma and Paraganglioma(PCPG) (AQ-AR), Prostate adenocarcinoma(PRAD) (AS-AT), Rectum adenocarcinoma(READ) (AU-AV), Sarcoma(SARC) (AW-AX), Skin Cutaneous Melanoma(SKCM) (AY-AZ), Stomach adenocarcinoma(STAD) (BA-BB), Testicular Germ Cell Tumors(TGCT) (BC-BD), Thyroid carcinoma(THCA) (BE-BF), Thymoma(THYM) (BG-BH), Uterine Corpus Endometrial Carcinoma(UCEC) (BI-BJ), Uterine Carcinosarcoma(UCS) (BK-BL), Uveal Melanoma(UVM) (BM-BN).
