## Supplementary figures and images for "*HspA1B* Is a Prognostic Biomarker and Correlated With Immune Infiltrates in different subtypes of Breast Cancers"

### SI-Fig 2

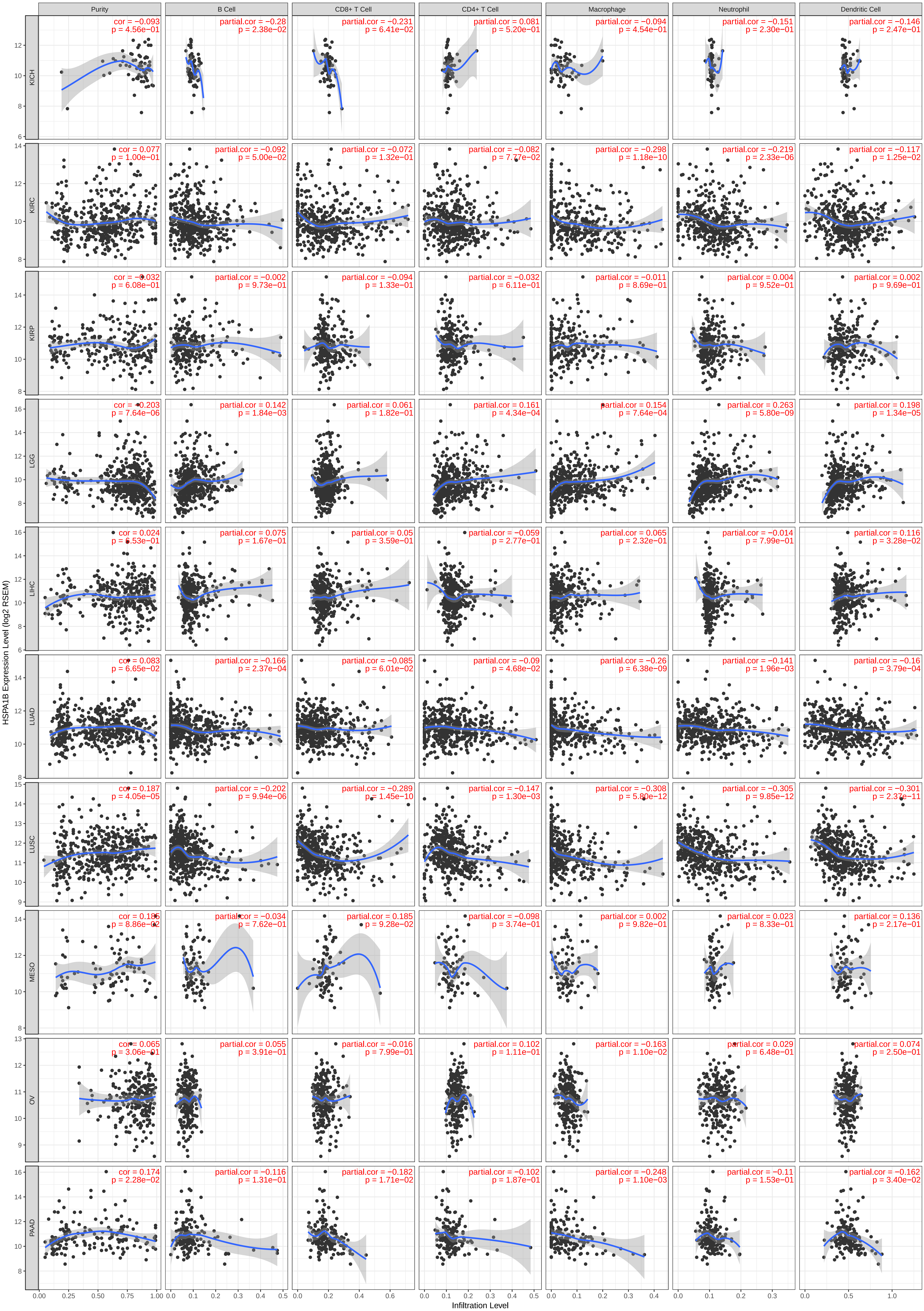

### SI-Fig 3

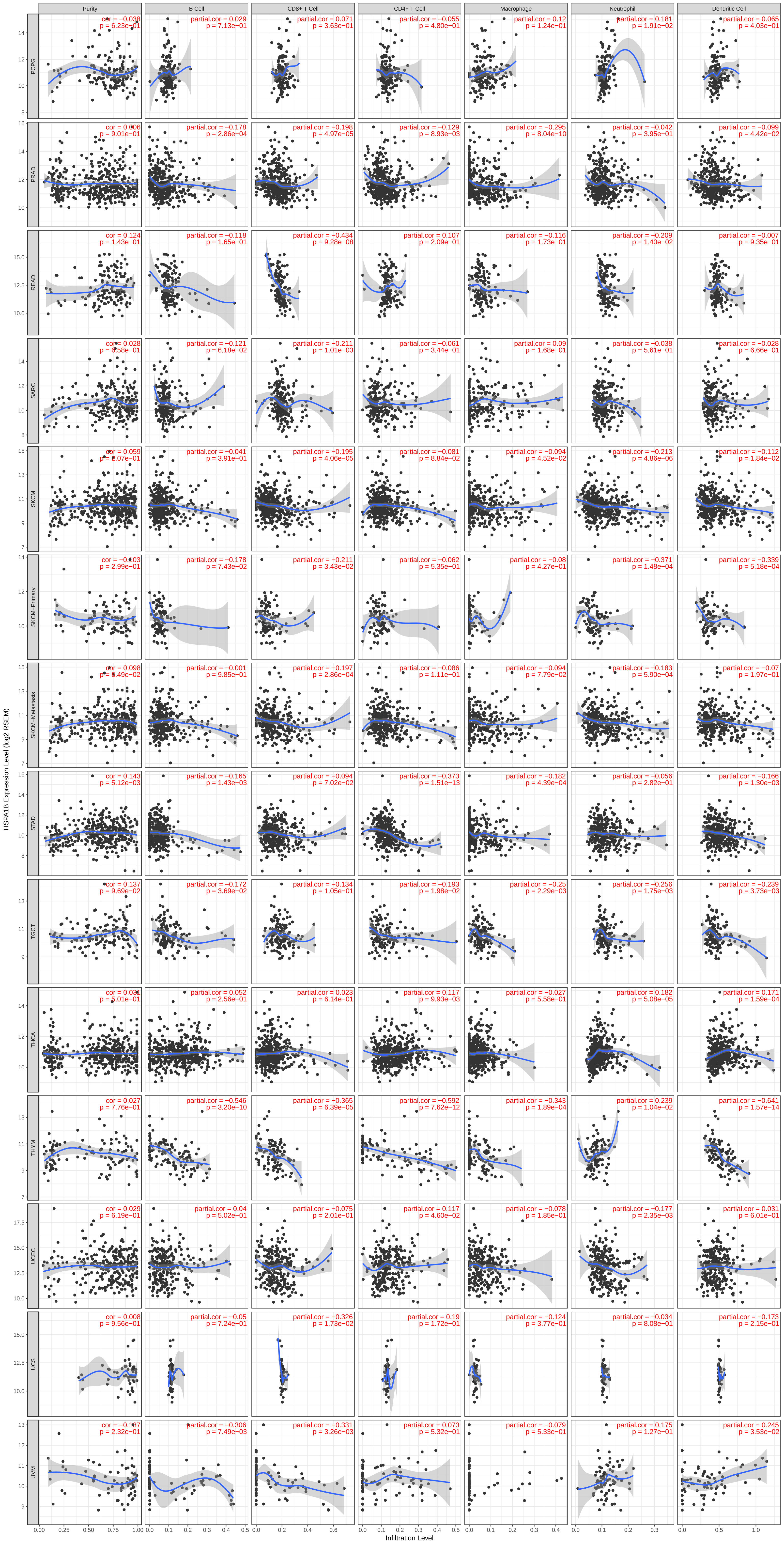

### Supplemental Data 1

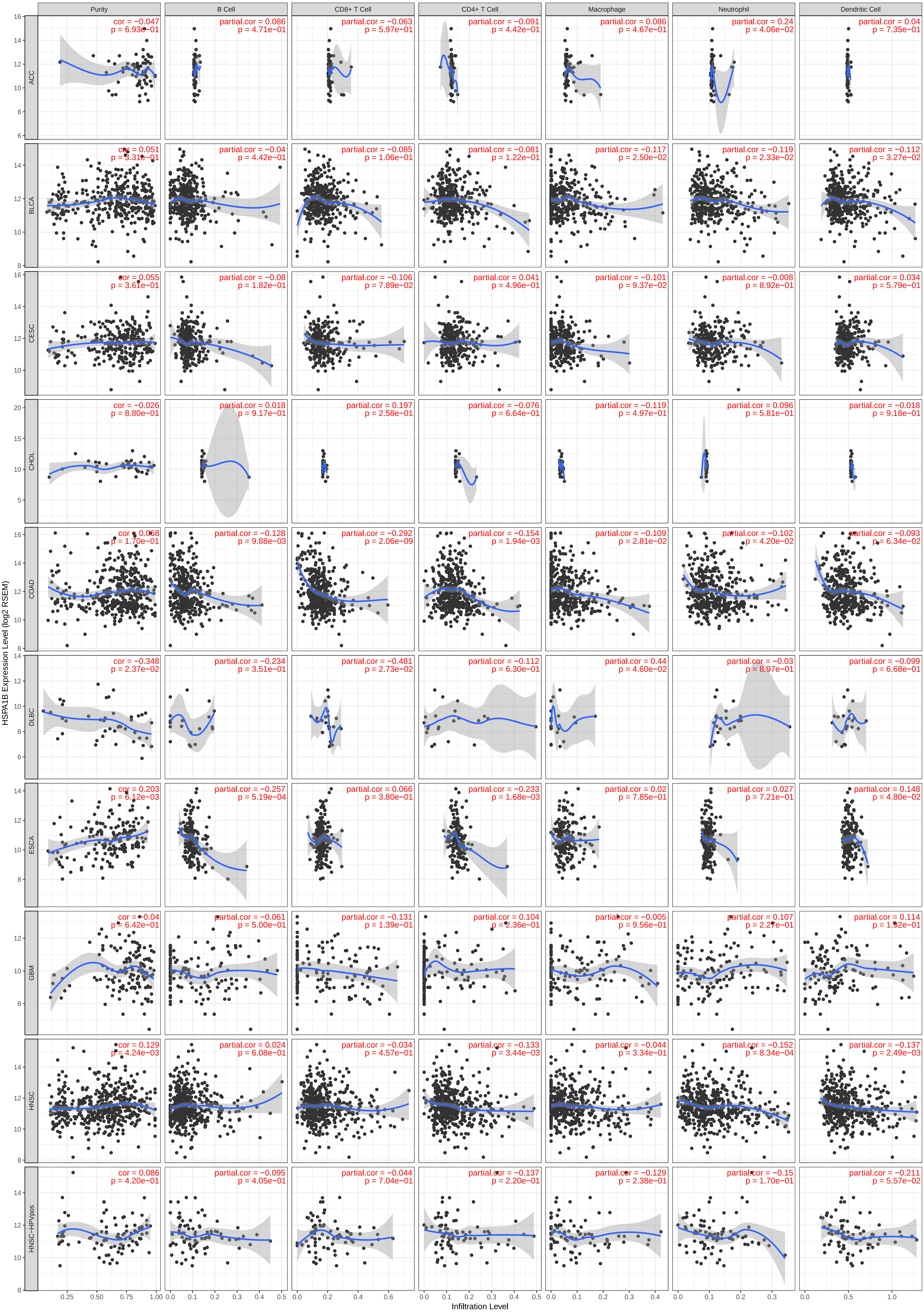
